# Spatial scale of transmission explains how acute infections circulate at low prevalence

**DOI:** 10.1101/2021.11.16.468880

**Authors:** Rebecca Mancy, Malavika Rajeev, Ahmed Lugelo, Kirstyn Brunker, Sarah Cleaveland, Elaine A. Ferguson, Karen Hotopp, Rudovick Kazwala, Matthias Magoto, Kristyna Rysava, Daniel T. Haydon, Katie Hampson

## Abstract

Fundamental questions remain about the regulation of acute pathogens in the absence of acquired immunity. This is especially true for canine rabies, a universally fatal zoonosis. From tracing rabies transmission in a population of 50,000 dogs in Tanzania between 2002-2016 we unravel the processes through which rabies is regulated and persists, fitting individual-based models to spatially-resolved data to investigate the mechanisms modulating transmission and the scale over which they operate. We find that while prevalence never exceeds 0.15%, we detect significant susceptible depletion at local scales commensurate with rabid dog movement, reducing transmission through clustering of rabies deaths and individuals incubating infection. Individual variation in rabid dog behaviour facilitates virus dispersal and co-circulation of lineages, enabling metapopulation persistence. These mechanisms likely operate in many pathogens circulating in spatially structured populations, with important implications for prediction and control, yet are unobservable unless the scale of host interactions is identified.

**One-Sentence Summary:** Identifying the spatial scale of contact reveals the mechanisms that limit the size of rabies outbreaks and that are critical for predicting transmission dynamics.

## Main Text

Understanding the mechanisms that regulate endemic disease prevalence remains a long-standing challenge in epidemiology (1, 2) and the mechanisms that enable long-term persistence at low endemic levels remain largely unexplored (3). This is particularly true for canine rabies, a fatal zoonotic virus for which naturally acquired immunity has not been demonstrated. The basic reproductive number R0 of rabies - that is, the expected number of secondary cases produced by a typical infectious individual in a fully susceptible population (4) - is low (between 1.1 and 2) and relatively insensitive to dog density (5), making the disease amenable to elimination through dog vaccination (6). Yet, dog-mediated rabies remains endemic across Africa and Asia where it kills tens of thousands of people every year (7) and its persistent circulation at such low prevalence in largely unvaccinated populations is an enduring enigma.

Rabies is, however, a uniquely tractable system for understanding how population-level patterns of infection emerge from pathogen transmission at the individual level. Rabies is transmitted via bites, which can often be observed, and the clinical signs are readily identifiable, with infected animals typically dying within one week of disease onset (fig. S1). Capitalising on these distinctive characteristics, we conducted exhaustive contact tracing to generate spatially resolved data on rabies infection and transmission in Serengeti district, Northern Tanzania, between January 2002 until December 2015. In this population of around 50,000 dogs (and 250,000 people) we traced 3612 rabies infections (Fig. 1, comprising 3081 cases in dogs, 75 in cats, 145 in wildlife and 311 in livestock), along with 6684 potential transmission events to other animals and 1462 people bitten by rabid animals of whom 44 died from rabies. Most cases could be statistically linked to progenitors, indicating that contact tracing detected the majority of cases. Further analysis suggested 83-95% case detection, with missed cases mainly being those generating limited, if any, onward transmission (Supplementary material). These data show that, in spite of local vaccination effort (coverage varying between 10 and 40%, fig. S3B), rabies circulated continuously with a maximum prevalence of just 0.1% of dogs.

**Fig. 1.**
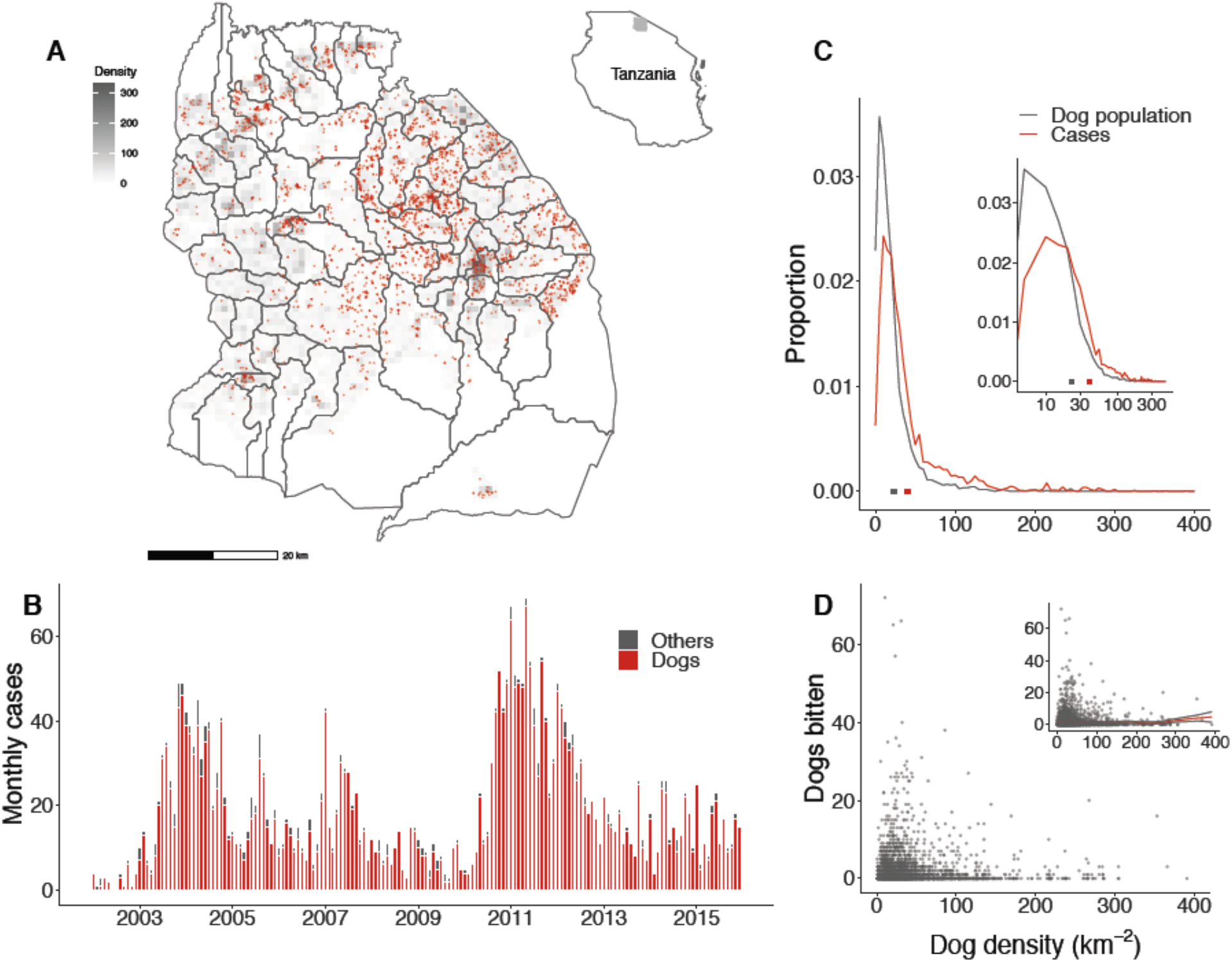
Rabies in Serengeti district from 2002 to 2016 in relation to dog density. **(A)** Case locations (red) mapped onto dog density (grey shading). Village boundaries are denoted by dark grey lines and the location of the district in Northwest Tanzania is shown in the inset. **(B)** Monthly time series of carnivore rabies cases (n=3,081) in domestic dogs (red) and other carnivores (grey, n=214; 75 cats, 60 jackals, 27 mongoose, 14 hyenas, 11 honey badger, 9 civet, 5 genet, 4 wild cat, 4 leopard, 3 serval and 2 foxes). **(C)** Proportional distribution of the dog population in relation to dog density (black, square shows mean of 23 dogs/km^2^) and of rabid dog case locations (red, square shows mean of 41 dogs/km^2^) in relation to dog density, indicating higher *per capita* incidence in higher density areas (independent samples t-test on log-transformed data, T=22.45, p<2.2e-16). Inset shows dog density and rabies cases on a logscale. **(D)** Dogs bitten per rabid dog, as determined by contact tracing, plotted against dog density at the location of each detected rabid dog (computed at the 1km^2^ scale). The inset shows a GAM fit to these data and the lack of an apparent relationship (p>0.05).

Endemic diseases are thought to be primarily regulated by the depletion of susceptible hosts, typically through disease-induced (or vaccine-acquired) immunity, counterbalanced by demographic processes (4). Yet the very low prevalence and absence of acquired immunity for rabies suggests that large-scale susceptible depletion must be negligible, challenging this explanation as a mechanism for persistence. We estimated dog densities across the district at high spatial resolution through a complete census, georeferencing almost 36,000 households and recording the vaccination status of dogs (in this setting almost all dogs are owned and free-roaming (8)). We saw no clear relationship between dog population density and contact rate when examining the numbers of dogs bitten per rabid dog (Fig. 1C), but mapping rabies infections revealed a small, yet significantly higher incidence of cases in areas with higher dog densities (Fig. 1C), suggestive of density-dependent processes. These observations are difficult to reconcile: how does transmission respond to dog population density and what processes keep prevalence so low?

Here, we propose that understanding the fine-scale structure of rabies transmission networks is critical to explaining its persistent dynamics and can inform its control and eventual elimination. We used the serial interval distribution - defined as the interval between the onset of infection in primary and secondary cases - and movement of traced rabid dogs to reconstruct local transmission trees. From these trees we identified putative introductions into the district and characterized clades of transmission descending from these index cases. Over the 14 years we estimated around 238 introductions (8-24 per year), most likely due to rabid dogs from neighbouring villages in adjacent districts (movie S1). These introductions led to locally sustained transmission, with 22 clades circulating for over 12 months (accounting for >70% of cases), and two clades each circulating for more than 4 years, illustrating how co-circulation of lineages contributes to persistence as part of a metapopulation (Fig. 2). We fitted an individual-based model, seeded by these introductions, to the spatially-resolved case and dog density data to investigate the processes modulating transmission, the scale over which they operate, and how they facilitate these metapopulation dynamics.

**Fig. 2.**
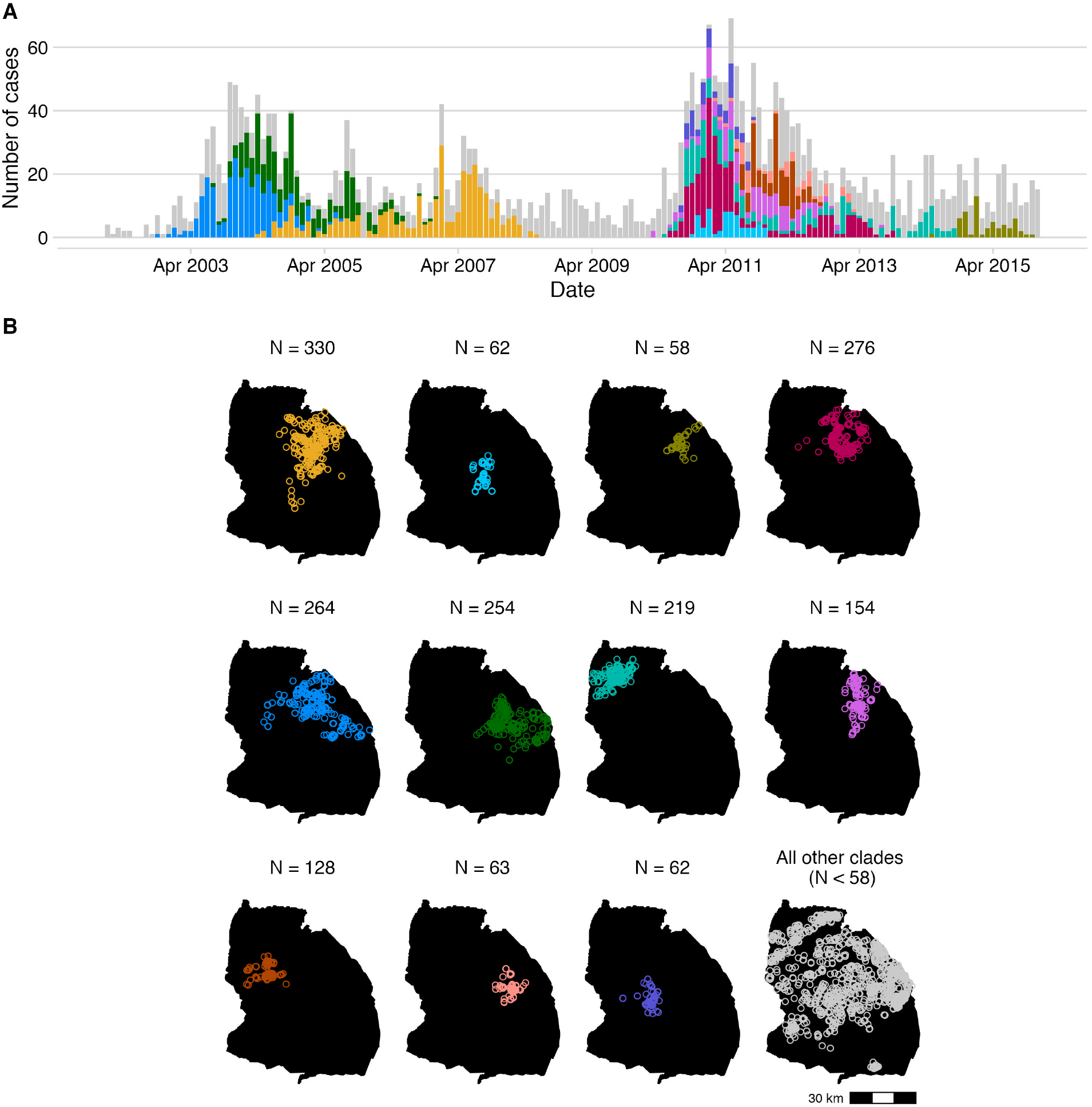
The spatiotemporal distribution of transmission clades. (**A**) Cases in each transmission clade per month (bars coloured by the 11 clades comprising the most cases, with all smaller clades shaded in grey) and (**B**) the spatial distribution of these largest clades, with the last panel showing the location of cases in all other clades (< 58 cases in size). The tree-building algorithm used the Lognormal serial interval distribution, the Weibull distance kernel and convolution and the 95% pruning threshold (Supplementary Materials).

We investigated the effect of host (dog) density on contact (biting) within our stochastic individual-based model using a Holling Type II functional response adapted from predator-prey theory (9). This functional response models contact rate as a saturating function of density, capturing a spectrum from weak to strong density dependence (10). By definition the contact process operates in accordance with population densities averaged over a particular spatial scale, but this scale is unknown. To understand the interplay with population density, we calculated contact rates according to the Holling Type II functional response at a number of different scales ranging from the whole district to spatially highly localised (within 0.25 km2 grid cells, fig. S2). From simulated transmission dynamics, allowing for dog population growth and varying susceptibility with demographic turnover and village-level dog vaccination campaigns, we used Approximate Bayesian Computation (ABC) to estimate the parameters governing the relationship between host density and contact rate together with individual variation in contact (biting). We designed the ABC acceptance criteria to capture temporal variability in rabies cases and their clustering by dog density (fig. S9) and determined the final posterior distribution and optimal spatial scale for modeling transmission according to how reliably accepted parameters reproduced observed dynamics.

Model fitting revealed that both the number of accepted parameter sets and their reliability in generating the observed dynamics differed according to the spatial scale of simulation (Fig. 3). Realistic dynamics were generated reliably only at the 1km^2^ scale, where the best parameter sets met the ABC criteria on >60% of re-simulations (fig. S10), accurately recreating case clustering and individual heterogeneity in contact (Fig. 3A and S1F inset). At other scales realistic dynamics were generated only rarely (fig. S10). At larger scales simulations in the absence of vaccination led to implausibly high disease incidence, yet at the 1km^2^ scale incidence remained generally realistic (Fig. 3B), with extremely rare runaway outbreaks (<1%) that human responses likely curtail in practice. We conclude that the processes that regulate rabies dynamics, including the size of outbreaks and overall prevalence, operate very locally, at scales typically much smaller than are modelled.

**Fig. 3.**
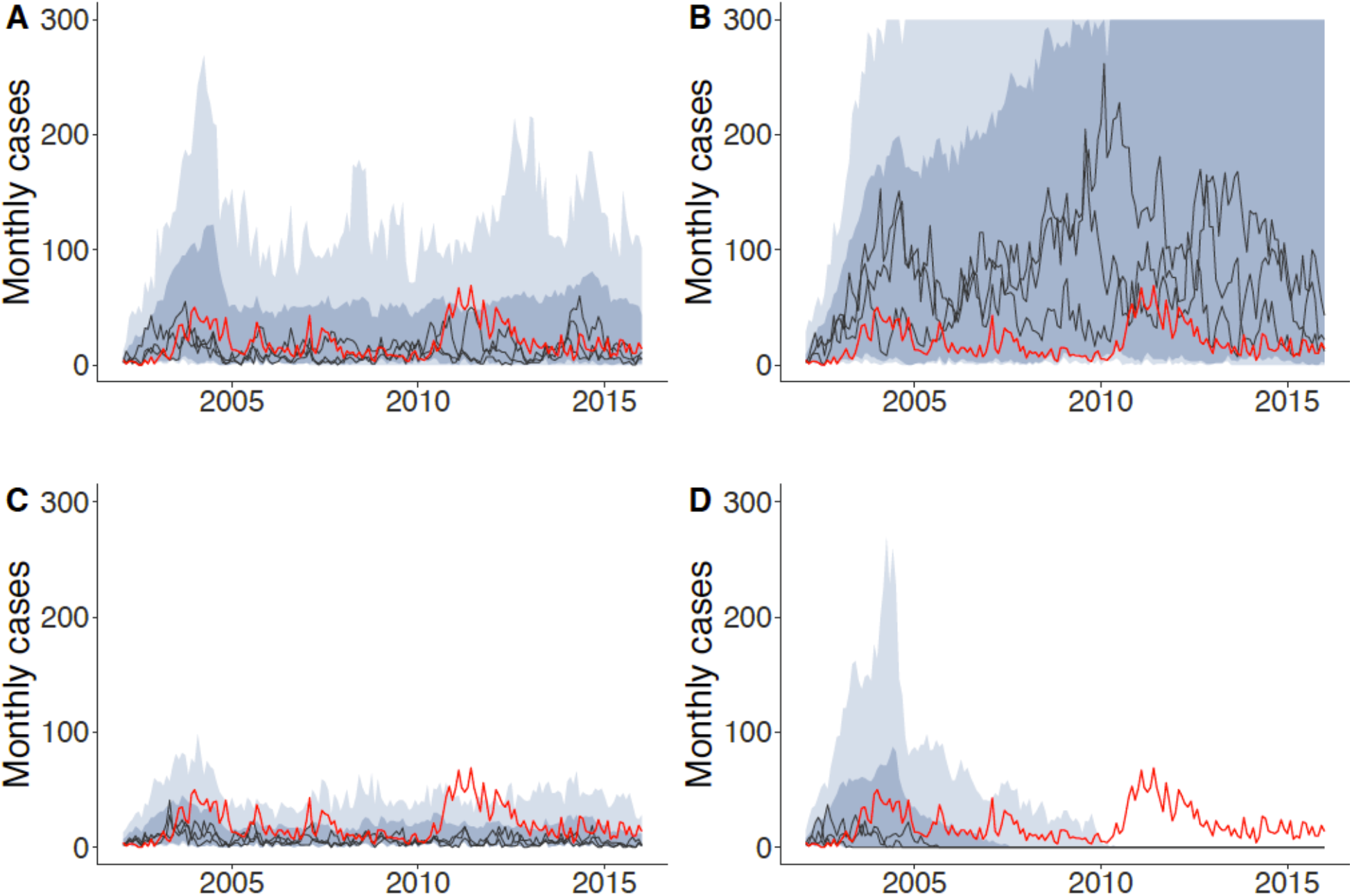
Comparison of observed time series of rabies cases and posterior simulations for different scenarios. Observed cases (red), plotted with interquartile range (dark shading) and 95% prediction intervals (light shading), both computed pointwise, from simulations at the 1km^2^ scale, and with 3 illustrative example runs (dark lines). The scenarios are simulated for (**A**) the final posterior distribution with vaccination campaigns as implemented; (**B**) under low levels of vaccination coverage; (**C**) with vaccination as implemented but no individual heterogeneity in contact parameters and (**D**) without incursions after the first year.

Simulating from the final posterior distribution (fig. S9) we decomposed the contribution of mechanisms regulating transmission, showing how susceptible depletion arises locally in two ways: from deaths of rabid dogs reducing contact opportunities, and through redundant exposures of dogs already incubating infection. The Holling curve illustrates how density-dependent contact rates at the lowest population densities asymptote to density independence at densities above the median (16 dogs/km^2^, Fig. 4). Simulations of index infections to estimate R_0_ (i.e. in an entirely susceptible population) show that rabid dogs bite, on average, 2.91 dogs leading to approximately 1.47 secondary cases per index case (95% Percentile Intervals (PI) 1.39-1.56). This R_0_ value is slightly higher than previously estimated (5), in part due to population growth (median dog density increased from 12 to >20 dogs/km^2^ over the 14 years) but varies across the landscape in relation to dog densities (Fig. 4) and according to how it is measured. R_0_ estimates from randomly selected index locations were slightly lower (1.35, 95% PI 1.27-1.43) than from randomly selected index dogs (proportional to density), while index cases from the transmission network, i.e., cases resulting from endemic simulations under low levels of vaccination coverage, generated marginally higher estimates (1.48, 95% PI 1.38-1.58), as they are slightly more prevalent in higher densities cells (Fig. 4). Under endemic circulation, local susceptible depletion reduces the effective reproduction number, R, by just over 30% (to around 1), of which 78% is from recent rabies deaths reducing contacts and 22% from reexposures of already incubating dogs. Thus, infected dog movement (fig. S1) determines the scale of susceptible depletion mechanisms that regulate endemic dynamics, such that even small outbreaks (~5 cases within 1km^2^) can substantively reduce R, given the heterogeneous distribution of dogs on the landscape (Fig. 4). There remains ambiguity in the mechanisms underpinning density independent transmission at higher dog densities. Human behavioural responses likely play a role (45% of traced rabid dogs were either killed or tied), and are to some extent captured in our model, but may operate differently during larger outbreaks (beyond those observed) and in more urbanized populations (<2% of dogs in this rural district live at densities >100 dogs/km^2^) and merit investigation.

**Fig. 4.**
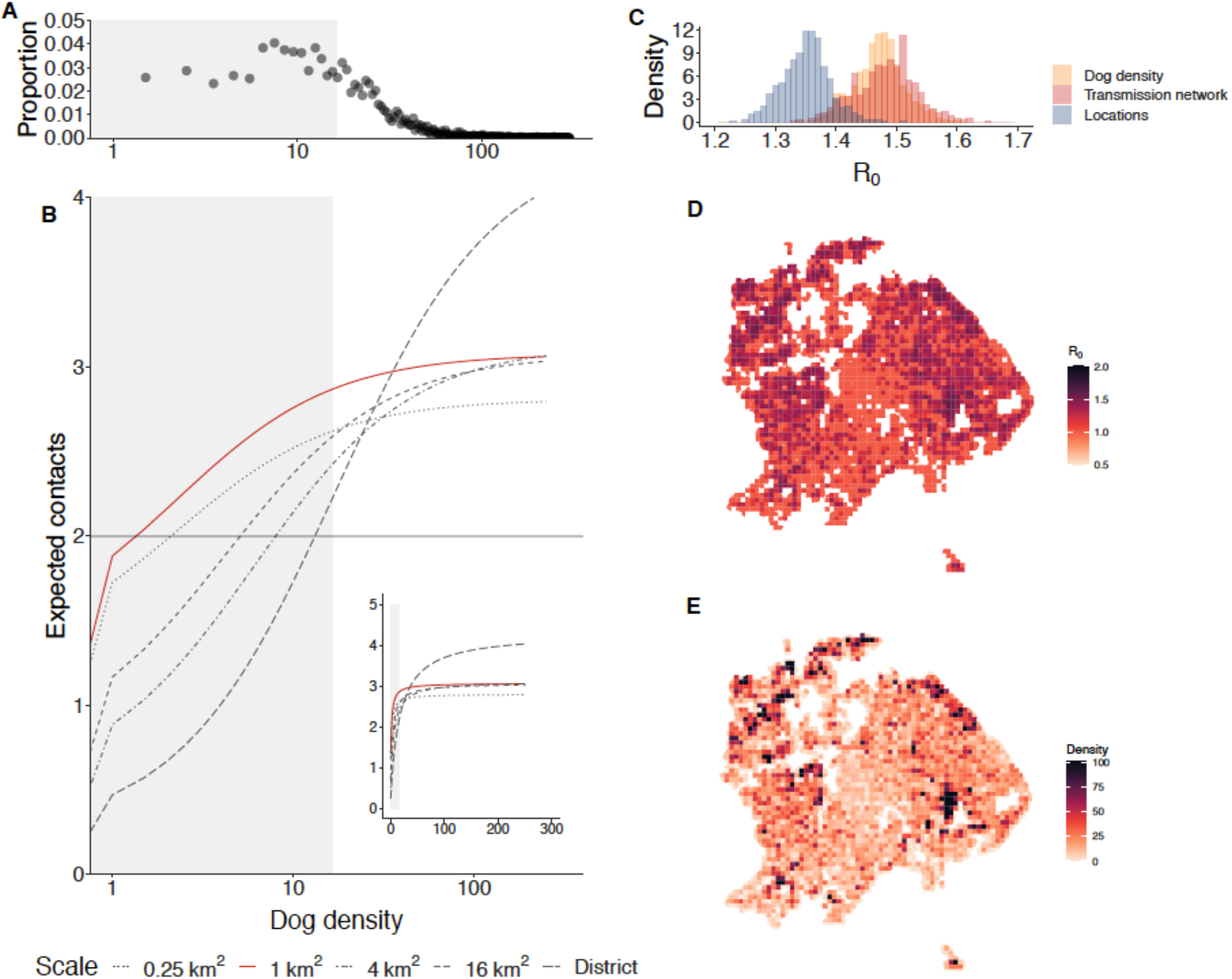
Rabies transmission in relation to population density. (**A**) Distribution of dog densities in Serengeti district shown on a log scale, with grey shading highlighting densities below the median (16.4 dogs/km^2^) midway through the time period (density increased from 12 to 22 dogs/km^2^ over the 14 years). (**B**) Holling curves computed from the most reliable parameter set at each scale, with the contact rate rescaled by the median infectious period (2 days) and dog density on a log scale; grey shading indicates dog densities below the median (16 dogs per km^2^) and the inset shows the holling curve by dog density on a linear scale. At the optimal spatial scale (1km^2^, red line) contact rates are density dependent at low dog densities and become increasingly density independent at higher densities. Parameter sets at other scales failed to reliably generate observed rabies dynamics. The Holling curve indicates how even the removal of a small number of dogs locally (from rabies deaths or incubating infection) reduces transmission such that R approaches 1 (corresponding to around 2 contacts per infection, of which around half develop rabies, horizontal grey line). (**C**) Histogram of R_0_ estimates from different approaches to simulating index infections, either by location (blue), density (orange) or from cases on the transmission network under low levels of vaccination coverage (red). (**D**) Mapped R_0_ estimated from simulating index infections sampled by dog density and (**E**) Mapped dog density at the midpoint of the time period.

Individual variation in transmission rate has been associated with rarer but more explosive outbreaks and more frequent extinction (14). We observed considerable variation in rabid dog behaviour (fig. S1), with a few rabid dogs biting many others (4% of rabid dogs bit >10 other dogs each and 4 bit >50 dogs) and traversing long distances (9 rabid dogs ran >10 km to bite other animals). The overdispersed distribution of clade sizes from our transmission trees (fig. S11), was in line with this variation in rabid dog biting behaviour, which was captured by our model-fitting approach (fig. S1F) revealing its importance in rabies dynamics. In counterfactual simulations from the final posterior distribution without individual heterogeneity the incidence of rabies was reduced by around 50% (fig. S7C) and the (relatively) large outbreaks observed in nature and in simulations with this heterogeneity did not occur (Fig. 3C versus A). The biological drivers of variation in contact result predominantly from individual-level (14), rather than environmental or population-level differences (15, 16) such as host density, but are poorly understood. Behavioural manifestations of infection that depend on sociological and pathological factors, like exposure dose and sites of viral proliferation (17), might underpin this individuallevel variation.

Both the scale of and heterogeneity in contact are crucial to capturing rabies dynamics (Fig. 3). Contact that leads to transmission includes density-dependent processes which well described theoretically (2, 17), are extremely challenging to quantify in relation to spatial scale, and this may explain why empirical studies of directly-transmitted diseases have not often found evidence of strong density-dependent transmission (18–20). For rabies, the main susceptible depletion mechanisms - deaths of rabid dogs and re-exposures of dogs already incubating infection - only curtail transmission if infection is clustered and explain why non-spatial models of rabies do so poorly at replicating dynamics, since rabies incidence is negligible at the population-level. Clustering has previously been shown to reduce transmission via the local build-up of immune individuals (13) and in the context of redundant biting by insect vectors (21); it has also been theorized to play a role in reducing transmission in the early stages of epidemics (22). For rabies, the incubation period acts in a similar way to immunity, resulting in redundant exposures that limit transmission. Natural immunity is not generally considered important in canine rabies or required to explain persistence, but antibodies have been detected in healthy unvaccinated dogs (11). If short-lived immunity does follow aborted infections, as thought for vampire bat rabies (12), our expectation is that it would cluster like incubating infections, reinforcing local scale effects. Although local susceptible depletion due to clustering has been shown to lead to less explosive epidemics and longer persistence times (18), the potential relevance of local susceptible depletion mechanisms to many other pathogens may be underestimated because their measurement relies on sufficiently resolved datasets. Our conclusion, that the relevant spatial scale at which to consider host density is determined by the scale of movements of infectious hosts, offers an avenue to interrogate the dynamics of pathogens for which spatially detailed data are lacking.

Our analyses further illustrate the degree to which introduced cases contribute to rabies persistence. In the absence of introductions and under observed levels of vaccination, we expect infection to circulate in the Serengeti district for up to 7 years (Fig. 3D). But, with between 8-24 rabid dogs introduced each year from neighbouring villages, infection persists even under reasonable vaccination coverage, despite most introductions causing only short-lived chains of transmission. In settings where vaccination coverage is negligible (i.e., dog populations across much of Africa and Asia) our simulations indicated a mean duration of outbreaks from single introductions of between 10-30 weeks, but the maximum exceeded 12 years (fig. S11). Locally self-limiting clusters of cases recur on the landscape (Supplementary movie), and in combination with heterogeneous movement and contact, permit the invasion and co-circulation of multiple lineages (19) (Fig. 2). Recurrent introductions and extinctions have been reported in many endemic areas (20–22) and cross-border introductions have led to rabies emergence in several previously rabies-free areas (23–27). In contrast to diseases like dengue (28), chains of infection circulate largely independently, given the exceedingly low prevalence of cases and only very spatially localised susceptible depletion. The concurrent extinction of all lineages therefore becomes less probable as more chains of infection co-circulate.

From a practical perspective, our findings explain why culling has typically proven so ineffective for controlling rabies, since dog populations would need reducing below very low densities across all areas where infection is circulating. Indeed, culling over 50% of the 400,000 dogs in Flores, Indonesia had no apparent impact on rabies circulation (29). Our results reinforce the message that mass dog vaccination remains the most effective and feasible method of controlling rabies, and provides insights that should inform elimination strategies. Although vaccination coverage varied in Serengeti district due to operational constraints, our simulations predict that on average rabies prevalence would have been at least three times higher under the low levels of vaccination typical in this region (Fig. 3B), and that the dog vaccinations in the district prevented at least 4,000 animal cases, 2000 human exposures and 50 deaths, although we do not consider how wildlife increases the susceptible population, or human responses during large outbreaks. Yet, dog vaccination has not locally eliminated rabies. This is both because the vaccination campaigns were not sufficiently comprehensive and introductions from neighbouring populations continually seeded new foci (Fig. 2). Coordinated scaling up of dog vaccination, as advocated by the global strategy to eliminate human deaths from dog-mediated rabies by 2030 (30), should therefore be expected to minimize introductions within connected landscapes and accelerate progress towards elimination. Moreover, sufficiently wide cordon sanitaire vaccination together with enhanced surveillance could keep areas rabies-free following local elimination, otherwise introductions from endemic areas will remain a threat.

Our results suggest that the predictive ability of epidemiological models for rabies, and perhaps for the endemic circulation of many other directly-transmitted pathogens that persist at low prevalence, such as Ebola in human (or ape) populations, crucially depend on whether they capture the effects of spatial and social/ population structure on susceptible depletion. For rabies, the local nature of interactions is missed by nonspatial models in contrast to our individual-based model. A further question is how these metapopulation dynamics scale up, and whether they can result in observed levels of large-scale synchrony (31). Our work also challenges the degree to which threshold quantities operate (32), for example, densities below which the virus cannot persist or vaccination coverage levels that rapidly lead to elimination. The clustered nature of transmission means that rabies can circulate locally for sufficiently long periods, enabling resurgence as vaccination coverage wanes, facilitated by metapopulation dynamics across heterogeneously distributed populations. Thus, in practice it is difficult to both determine and apply such threshold concepts and future theoretical work is warranted. A further difficulty for evaluating how accurately models predict the impact of rabies control measures is limited data. Surveillance for a zoonotic pathogen that circulates at such low prevalence is challenging, especially since routine surveillance is weak in countries with endemic canine rabies. Work is therefore needed to strengthen surveillance and to assess case detection, while models need to be developed for use with routine surveillance to assess realistic time horizons for elimination and the scales over which control programmes are best implemented. Concurrent circulation of viral lineages offers an opportunity for using genomic data to assess the performance of both rabies surveillance and control, particularly for differentiating undetected circulation from reintroductions (33).

Heterogeneous mixing fundamentally changes infection dynamics (13) but is not well understood in relation to the mechanisms underpinning the endemic circulation of pathogens in nature. Our study captures key ways in which individual variation in contact and movement, and their interaction with host density, play out at the population level. Though often cited as being important for understanding the relationship between transmission and host density (34, 35), spatial scale is rarely investigated. Our uniquely detailed and long-term data on rabies transmission in conjunction with the underlying population distribution allowed us to jointly estimate individual variation in contact and the spatial scale of interactions, and thus identify crucial mechanisms that regulate rabies dynamics and that explain how the disease can circulate at such low prevalence. Specifically, we show that localized susceptible depletion mutes rabies outbreaks, whilst infrequent longer distance movements by rabid dogs’ seed cases in unaffected localities away from the effects of local susceptible depletion, therefore playing a disproportionate role in facilitating persistence.

## Acknowledgments

We thank local communities and government staff from the animal and public health sectors for ongoing support, and the Serengeti Health Initiative for continued dog vaccination support. Jessica Metcalf, Daniel Streicker, Jonathan Dushoff, Michael Li and three reviewers provided very constructive feedback, which greatly improved the work. The study was approved by the ethical review board of the Ifakara Health Institute and the National Institute for Medical research in Tanzania. We are grateful to MSD Animal Health for donating vaccines for dog vaccination campaigns.

## Author contributions

Conceptualization: KH, RM, DTH

Methodology: RM, MR, EAF, KH

Investigation: KH, MM, AL, KR, MR, KB, KHo

Visualization: KH, RM, MR, EAF

Funding acquisition: KH

Project administration: KH, RK

Supervision: KH, RK

Writing – original draft: KH, RM

Writing – review & editing: KH, RM, DTH, SC, KR, MR

## Data and materials availability

Anonymised data and code to reproduce the analyses are available from Github: *https://github.com/boydorr/RabiesTransmissionScale*. The contact tracing and the census involved collection of personally identifiable information. Individuals interested in accessing these data should contact KH to organize ethical clearance.

## Supplementary Materials

Materials and Methods

Figs. S1 to S13

Tables S1 to S2

References (*36*–*40*)

Movie S1

