## Supplementary Materials for "Spatial scale of transmission explains how acute infections circulate at low prevalence"

**Materials and Methods**

Overview

We collected three datasets comprising: 1) probable animal rabies case histories from contact tracing; 2) spatially-resolved dog population data from a georeferenced census conducted throughout the Serengeti district; and 3) dog vaccination information compiled from campaign records. We used the contact tracing data to directly estimate key epidemiological parameters which we used in the construction of transmission trees. We developed a simulation platform for modelling rabies transmission, dog demography and dog vaccination campaigns in order to estimate three contact parameters. These parameters were derived from the Holling functional response and allowed us to determine the functional form of contact in relation to population density.

We initialized our model using the introductions identified from our transmission tree reconstructions and processed the contact tracing datasets for model fitting using Approximate Bayesian Computation (ABC). For the ABC, we designed acceptance criteria to capture the spatial and temporal variability in the occurrence of rabies cases and biologically plausible limits to transmission. We used two rounds of ABC to define a final posterior distribution and the best spatial scale for modeling rabies dynamics. We subsequently compared the contact tracing data with several counterfactual scenarios that we explored by simulating from the final posterior distribution, as well as decomposing the processes regulating transmission, by comparing simulations from index cases and from endemic scenarios. Finally we undertook a range of sensitivity analyses of our transmission tree reconstructions and comparisons with simulations to estimate the proportion of cases detected using contact tracing and to understand how incomplete case detection impacts our findings.

The study was approved by the Tanzanian National Institute for Medical Research (NIMR/HQ/R.8a/vol.IX/2788), the Ministry of Regional Administration and Local Government (AB.81/288/01) and Ifakara Health Institute Institutional Review Board (IHI/IRB/No:22-2014).

Data collection

*Probable animal rabies case histories*

Probable animal rabies case histories were obtained through exhaustive contact tracing, as described in (1, 2). These data consisted of a georeferenced record for each animal known to have either been bitten by a probable or confirmed rabid animal, or to have shown signs of rabies. Each record included the date when signs of rabies were first observed; where known, a unique identifier of the infectious animal that bit it and the date when this occurred; and the outcome of the animal. A total of 3,619 probable animal rabies cases were recorded from 2002-2015 (Figure 1), including 3,081 cases in dogs, 311 in livestock, 75 in cats and 145 in wildlife (mostly jackals, mongoose and hyena species as well as honey badgers, civets, genets, serval, wild cats, leopards, bat-eared foxes, a baboon and a wildebeest). Of these cases, 348 were confirmed through diagnostic testing. A further 5,414 animals were recorded to have been bitten by probable rabid animals during this period but not to have developed rabies. Of the 3,281 rabid carnivores found via contact tracing, we identified progenitors for 1148 (35%) and recorded an incubation period for 1,153 (35%). Of these, 397 (12% of the total) had information on the infectious period (Figure S1).

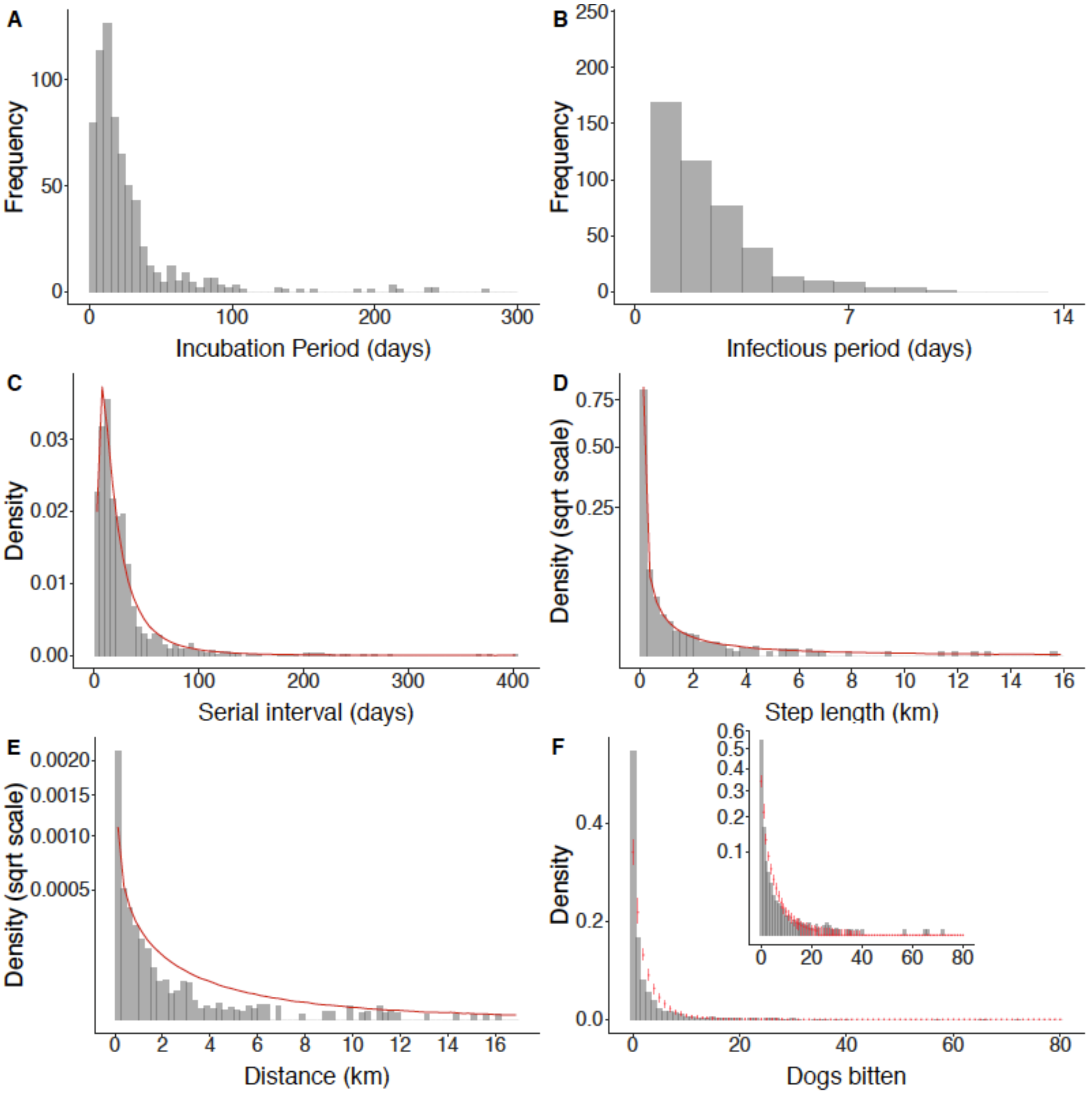


**Fig. S1. Attributes of rabid dogs measured from contact tracing.** Empirical distributions of: (A) Incubation and (B) Infectious periods; (C) Serial intervals; (D) Step lengths inferred between bites made by rabid dogs (see S3); and (E) Distances between biters and their contacts. The best-fitting distributions to these data, that were used either in the transmission tree reconstructions or the simulations are shown in red (further details in Table S1). Panel (F) shows the distribution of dogs bitten per rabid dog as determined through contact tracing (grey bars), with the inset plotted with the y-axis square root transformed. The simulated distribution of dogs bitten per rabid dog from the final posterior parameter sets at the 1km$^2$ spatial scale are overlaid as red lines (5th to 95th percentile). Note that the distances and step lengths were treated as censored for animals where their starting locations were unknown.

*Dog population density and distribution*

A complete human and dog population census was conducted from April 2008 until March 2016, recording the number of adults, children, dogs and puppies (<3 months old) per household, the rabies vaccination status of dogs and a household GPS location. From the census we counted 35,947 households, with 255,137 persons, 63,005 dogs and 22,062 cats (Figure 1). We combined counts of adult dogs and puppies to calculate the dog population per 0.25km2 grid cell on the date of the census. Populations per grid cell for each month were projected forward and back using an exponential growth model, applying human population growth rates derived from village-level population sizes (~2.5% per year) from the National Population and Housing Census (3). This allowed us to map the projected dog population over time, which increased from around 43,000 dogs at the start of 2002 to around 71,000 by late 2015. Monthly projected dog populations were aggregated per grid cell at five spatial scales: 0.25km2, 1km2, 4km2, 16km2, and for the entire district (i.e. fully mixed). Mapped population densities at the midpoint of the period (1 January 2009), aggregated at each spatial scale, are shown in Figure S2.

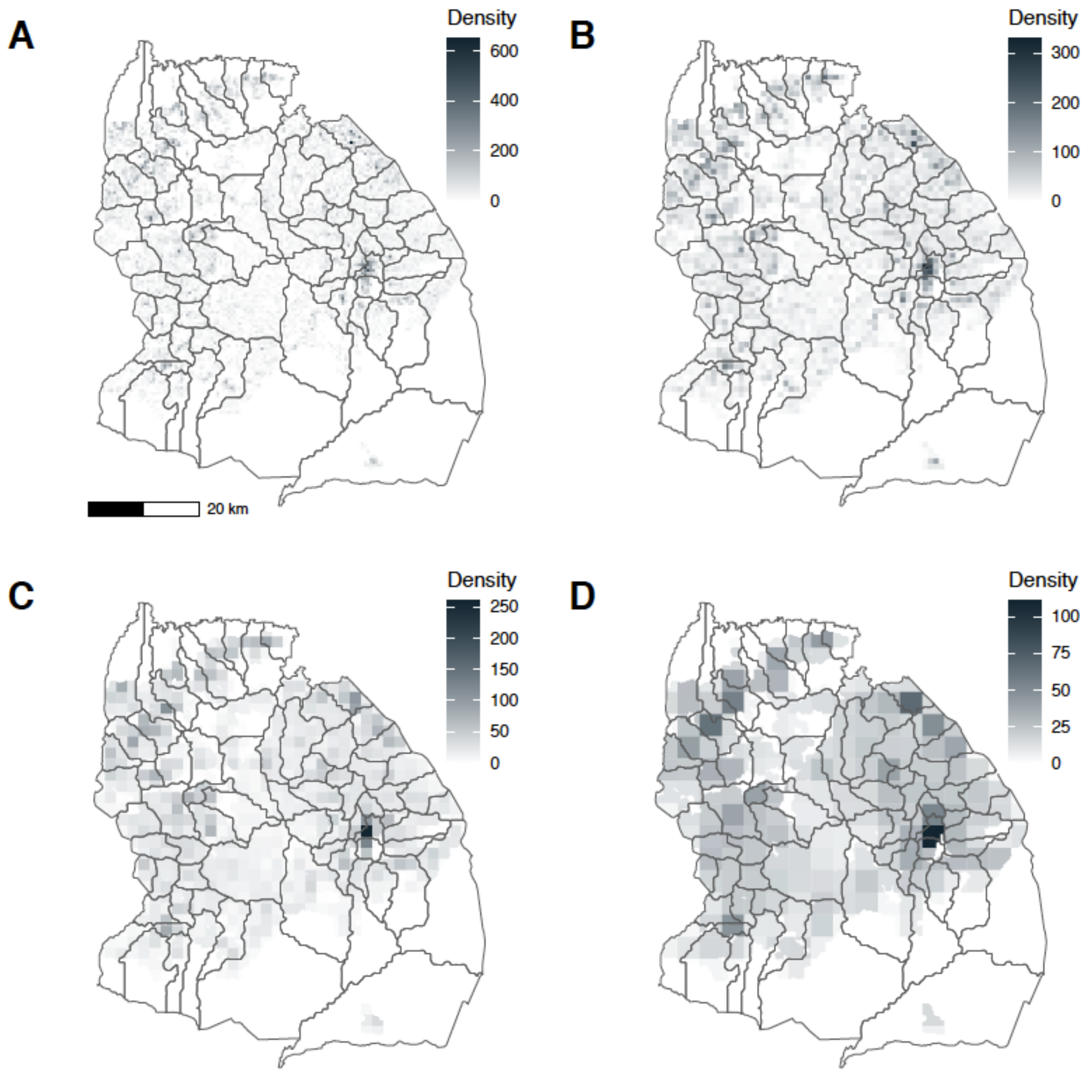


**Fig. S2. Dog densities aggregated at different spatial scales.** Dog densities are shown per $km^2$ at the midpoint of the simulation period, with 2002 village boundaries overlaid. (A) $0.25km^2$; (B) $1km^2$; (C) $4km^2$; (D) $16km^2$. On 1 January 2009, the average population density in the district was 23 dogs/$km^2$, with the highest density in Mugumu (the district town centre), reaching 330 dogs/$km^2$, measured at the $1km^2$ scale. Note the increasing density legend with the increasingly resolved spatial scale.

*Vaccination campaigns*

Records of vaccination campaigns at village or sub-village level were compiled, including the date of each campaign and the number of dogs and cats vaccinated. Campaigns were conducted in all years, but varied in timing and the number of villages reached due to logistic constraints (Figure S3). The number of dogs vaccinated each year generally increased over the period (Figure S3A), as did the proportion of total dogs that were vaccinated (Figure S3B). Highest coverage was typically achieved in areas of high dog density, which were most accessible for vaccination.

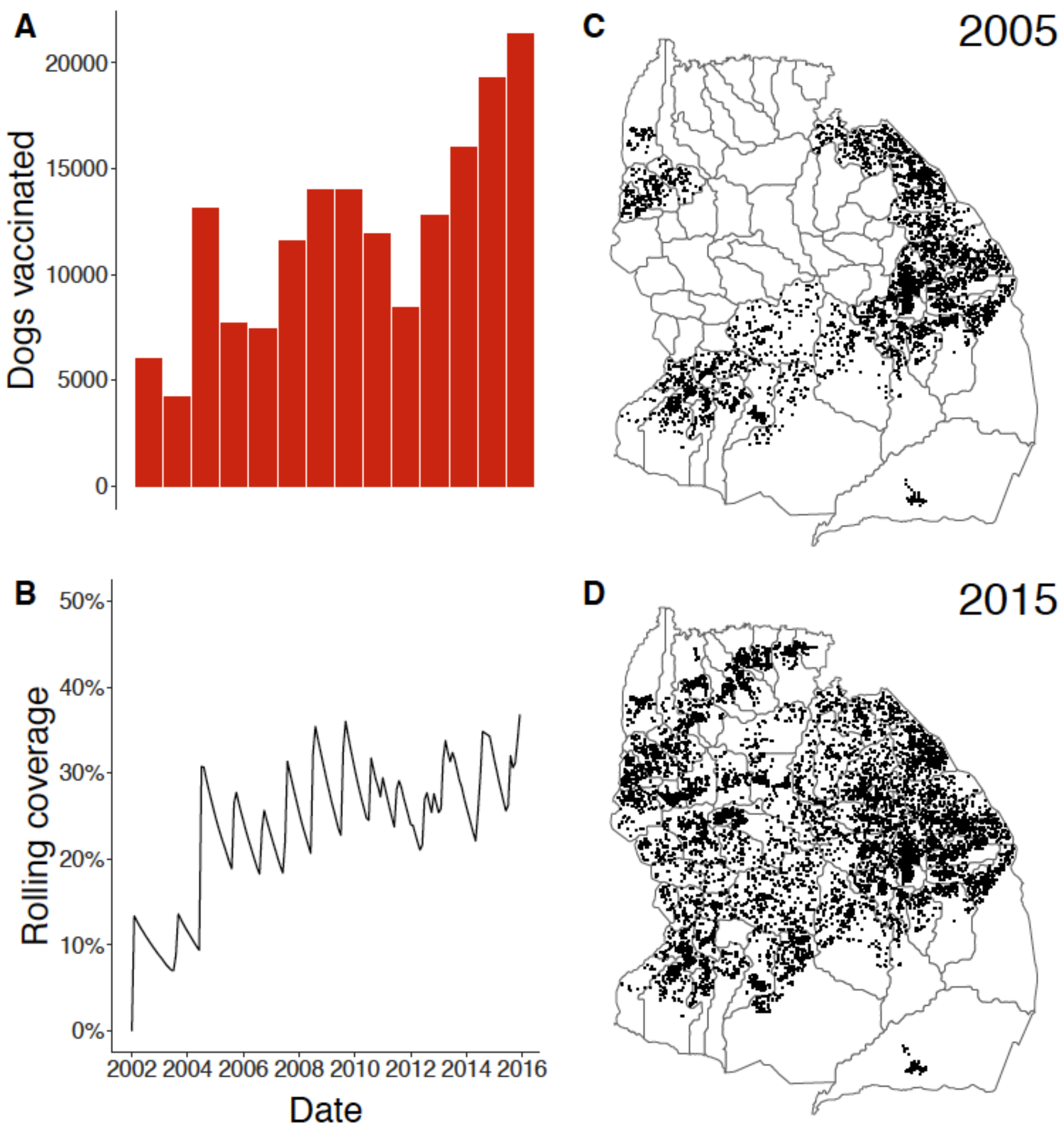


**Fig. S3. Dogs vaccinated each year and time-varying vaccination coverage.** (A) Total number of dogs vaccinated by year; (B) Time series of overall coverage at the district level from the simulations (vaccinations applied at the 1km$^2$ scale); Areas covered by vaccination campaigns in (C) a year with relatively poor spatial coverage and (D) a year with good spatial coverage. Black dots correspond to dog vaccinations from the village-level campaigns that year allocated according to our georeferenced census of dogs in the district.

<u>Epidemiological parameter estimation</u>

For the simulations, incubation and infectious periods were sampled directly from the data. Infectious periods were recorded as integer days. As periods shorter than one day were not quantified accurately, uniform random noise in the range (-0.5, 0.5) was added to infectious periods to avoid spurious discreteness. Dogs that were killed without biting often had no recorded infectious period, even when the incubation period was recorded. To retain realistic heterogeneity in incubation periods, we imputed infectious periods for 279 animals that were killed without biting, and for which incubation periods but not infectious periods were recorded. For this purpose, a number of days for the infectious period was drawn from an exponential distribution to model a waiting time until being killed or restrained (mean 0.5 days to ensure low probability of this time exceeding the maximum realistic infectious period of 7 days). Thus, in total, 676 pairs of incubation and infectious periods were used in the simulations.

Movement of rabid animals was modelled as steps between sequential bites, and was inferred from the locations of cases and the animals that they were recorded to bite in the contact tracing data. Because it was not always possible to determine the order in which rabid animals bit other animals, we computed step lengths in metres between bite locations following estimation of the solution to the travelling salesperson problem, starting from the location of the biting animal and excluding return distance (4). All steps were combined (total 5388 steps; mean 198m; sd 664m), and Gamma, Lognormal and Weibull distributions were fitted by maximum likelihood to the distribution of step lengths, left-censored at 100m to account for spurious zero step lengths when GPS locations were recorded at a homestead where multiple dogs were bitten. We selected the best-fitting distribution using the Akaike information criterion (AIC) (Figure S1C, Table S1) and used it in our simulation for drawing step length distances (The simulation platform).

We estimated distributions for the serial interval *S*, and distance kernel between cases *K*, respectively, for the purposes of constructing transmission trees (see below) and inferring introduced cases, i.e. cases that resulted from infection outside of Serengeti district, either from humans transporting a dog incubating infection into the district or from an infected dog transgressing from a neighbouring district. We retrieved 1,107 serial intervals from the data, each calculated as the interval between the dates symptoms started for a bitten case and its recorded biter. For the distance kernel, we computed the Euclidean distance between locations of cases and their traced contacts (including bitten but previously vaccinated dogs that did not develop rabies), and fitted distributions to these distances with a number of censoring assumptions. For dogs whose owner was known, and home location was therefore recorded accurately, we assumed left-censoring at 100m to account for zero distances recorded for animals biting at their own homestead. For animals of unknown origin (including wildlife), where the recorded location is the location the animal was first observed biting, rather than an accurate home location, we assume that recorded distances of <100m were actually at least 100m (as otherwise the animal would have been recognised) and that distances were right-censored. We fitted probability distributions (Gamma, Lognormal and Weibull) to the data on both the serial interval and the distance kernel.

The Lognormal, Gamma and Weibull distributions all fitted the serial interval and distance kernel data well, but the Lognormal and the Weibull distribution had the lowest AIC for these two distributions respectively (Table S1). Estimated parameter values are reported in Table S1, and the data and best-fitting distributions i.e. with the lowest AIC are shown in Figure S1.

| **Parameter** | **Median (95% range)** | **Distribution** | | **Value (CIs)** | | **Value (CIs)** | **AIC** | **N** |
|---|---|---|---|---|---|---|---|---|
| Incubation period | 20.096 (1.046-95.889) | Gamma | shape | 1.148 (1.064-1.231) | rate | 0.042 (0.038-0.045) | 9945 | 1153 |

| | | | | | | | | |
|---|---|---|---|---|---|---|---|---|
| (days) | **16.8 (2.552-110.618)** | **Lognormal** | **meanlog** | **2.821 (2.766-2.877)** | **sdlog** | **0.962 (0.922-1.001)** | **9692** | |
| | 18.927 (0.668-102.511) | Weibull | shape | 0.99 (0.95-1.029) | scale | 27.411 (25.713-29.109) | 9958 | |
| Infectious period | **2.37 (0.437-7.094)** | **Gamma** | **shape** | **2.427 (2.09-2.764)** | **rate** | **0.887 (0.759-1.016)** | **2084** | **588** |
| (days) | 2.292 (0.675-7.779) | Lognormal | meanlog | 0.83 (0.776-0.883) | sdlog | 0.623 (0.579-0.668) | 2095 | |
| | 2.396 (0.291-6.947) | Weibull | shape | 1.571 (1.46-1.682) | scale | 3.026 (2.857-3.196) | 2090 | |
| Serial Interval | 20.697 (1.074-98.853) | Gamma | shape | 1.146 (1.061-1.231) | rate | 0.04 (0.037-0.044) | 9614 | 1107 |
| (days) | **17.296 (2.604-114.877)** | **Lognormal** | **meanlog** | **2.85 (2.794-2.907)** | **sdlog** | **0.966 (0.926-1.006)** | **9380** | |
| | 19.466 (0.682-105.831) | Weibull | shape | 0.987 (0.947-1.028) | scale | 28.215 (26.428-30.003) | 9626 | |
| Distance Kernel | 797.012 (4.096-7159.644) | Gamma | shape | 0.589 (0.567-0.611) | rate | 0 (0-0) | 43769 | 6626* |
| (m) | 641.016 (21.641-18987.214) | Lognormal | meanlog | 6.463 (6.411-6.515) | sdlog | 1.729 (1.678-1.779) | 43905 | |
| | **747.536 (6.514-8205.369)** | **Weibull** | **shape** | **0.698 (0.678-0.717)** | **scale** | **1263.954 (1201.757-1326.15)** | **43734** | |
| Step length | 8.785 (0-1724.491) | Gamma | shape | 0.151 (0.142-0.161) | rate | 0.001 (0.001-0.001) | 28784 | 5388 |
| (m) | 31.714 (0.515-1954.018) | Lognormal | meanlog | 3.457 (3.355-3.558) | sdlog | 2.103 (2.015-2.19) | 28843 | |
| | **22.192 (0.004-1726.144)** | **Weibull** | **shape** | **0.384 (0.369-0.399)** | **scale** | **57.641 (51.833-63.45)** | **28757** | |

**Table S1.Estimated epidemiological parameters.** Confidence intervals for each parameter estimate are shown as well as the median and 95% quantile range of the data, the AIC and the number of data points for each estimate. For each parameter, the distribution with the lowest AIC is shown in bold. The serial interval and distance kernel were used for reconstructing transmission trees, and the step length distribution was used for simulations with incubation and infectious periods drawn directly from the data.*3,275 points were treated as right-censored due to the unknown start location of biter.

<u>Transmission tree reconstructions</u>

Using the contact tracing data, we reconstructed transmission trees probabilistically to infer transmission chains and assign possible introductions of rabies into the district. We adapted an algorithm to construct trees by assigning putative biters to each reported bitten case in rabid carnivores (i.e. domestic dogs, cats, or wildlife) as identified from contact tracing, or based on the spatiotemporal window between cases and the estimated distance kernel K and serial interval S distributions, accounting for uncertainties in timings and locations (1). For each case i, we selected a biter at random with probabilities pij from all j∈{1,...,n} cases preceding that case, where:

$$p_{ij} = \frac{S_{ij}K_{ij}}{\sum_{k=1}^{n} S_{ik}K_{ik}}$$

$S_{ij}$ is the probability of the time in days between case *i* and its potential biter *j,* where the time interval must exceed zero*;* and $K_{ij}$ is the distance between the locations of case *i* and biter *j*. Because the dates when some individuals were bitten or developed rabies were only approximately known, 1,000 bootstrapped datasets were generated with dates drawn randomly from a uniform distribution over the window of uncertainty. Within trees which incorporate known traced biter-bitten pairs but still recorded some uncertainty in precise dates, dates were drawn from the window of uncertainty for these cases, but constrained so that a known biter's date of symptoms is at least seven days before it's earliest known secondary case.

For probable cases in wildlife (n = 139) and for dogs and cats where the owner was not known (n = 1283), i.e. 1422/ 3295 cases, we used the convolution of *K* to account for the additional uncertainty in the recorded coordinates for these cases. To parameterize this distribution, we fitted a probability distribution to a simulation of two dispersal events with a direction drawn at random (uniformly from 0 to 360 degrees) between movements (i.e. from the location of the biter to the unknown animal, to the location of the unknown animal when it bit another animal or person, which was the georeferenced point for the animal recorded whilst contact tracing). Distances of <100m were censored to match the fitting used for the distance kernel distributions.

We applied a pruning approach to distinguish transmission clades and their index cases. Within the tree-building algorithm, cutoff values from the estimated serial interval and distance kernel distributions (S and K), were used to define plausible progenitors (5), except for known biters, where the distance or time between cases was ignored. From 1000 bootstrapped trees we generated a consensus tree as the set of biters most frequently selected for each case. We designated introductions as those cases which seeded a transmission clade in the consensus tree (i.e. cases for which no biter was assigned given the pruning cutoff). Given date uncertainties, there can be ambiguity in who-infected-whom, which generates loops within the consensus tree (i.e. case i is most often selected as the biter for case j, and case j is most often selected as the biter for case i). To break these loops, we selected the next most likely biter for the case with the earlier date of symptoms in the case pair. We did this sequentially for each looped pair, so that no new loops could be generated. If a case had no other selected biters within the set of bootstrapped trees, it was assigned as an introduction. The tree-building methods were wrapped into an R package, available at *github.com/mrajeev08/treerabid* and archived on Zenodo (*doi: 10.5281/zenodo.5269062*).

To parameterize the modelling, we reconstructed trees using the best fitting serial interval and distance kernel distributions, i.e. Lognormal for *S* and Weibull for *K* (Table S1, Figure S1), with the convolution of the distance kernel distribution for probable cases in wildlife or for dogs where the owner was not known. Our sensitivity analyses (see below) showed that these distributions better captured the long tails in the contact tracing data while mediating trade offs between identifying directly linked cases (high agreement with contact tracing) and introductions (low number mispecified). Traced transmission links were incorporated as known and biters assigned probabilistically for cases without a traced biter. Given that contact tracing was unlikely to detect all introductions into the district (see *Estimating case detection*), we applied a less conservative 95% pruning threshold within the tree-building algorithm to define introductions.

The consensus tree of inferred transmissions and introductions (n = 238, i.e. an average of 17 and a range of 8 - 24 introduced cases per year) is illustrated in the supplementary movie (movie S1). We highlight how these introduced cases led to sustained transmission within the district in Figure 2, where we colour the 11 largest clades from the consensus tree, which each contained >58 cases.

**Movie S1. Rabies cases and inferred transmission within Serengeti District from January 2002 to December 2015.** Transmission tree reconstruction using the consensus links from the tree-building algorithm with a Lognormal serial interval, Weibull distance kernel and convolution and 95% pruning threshold. Cases are animated each month, with animals that are incubating infections shown as empty circles until infectious when they transition to filled circles (note that many cases become infectious within the same month of exposure). Inferred transmission links are shown by curved lines at the approximate time of the exposure event, with colors of cases and lines corresponding to the transmission clades. The 11 largest clades are each plotted with a unique color, while all others are shown in grey. Cases identified as introductions are designated by a filled square. The top panel shows the monthly time series of cases by transmission clade.

Available here: https://www.dropbox.com/s/34uj01tfrujijki/animation.mp4?dl=0

*Sensitivity analyses of tree reconstructions*

We examined how sensitive transmission trees were to the choice of serial interval and dispersal kernel distributions used, noting that the Lognormal distribution allows more rare events i.e. long incubation periods or longer distances to secondary cases; events that we expect to not be as well captured by contact tracing (Figure S1). For each of these distributions we compared pruning cutoffs at 95% and 97.5% (Table S2) and examined the resulting range of introduction estimates from the consensus trees. To compare the performance of the tree-building algorithm, we also reconstructed trees without using traced transmission links and evaluated agreement between the consensus tree and the contact tracing data for the 1148 (35%) rabid carnivores with identified biters. We evaluated both the proportion of these cases for which biters were correctly assigned and the number which were misidentified as introductions.

| Probability distribution | Pruning threshold (%) | Distance cutoff in meters for animals with known vs unknown owners and wildlife (convolution) | Time cutoff in days |
|---|---|---|---|
| Gamma | 95.0 | 2696 (8300) | 81 |
| | 97.5 | 3471 (10165) | 99 |
| Lognormal | 95.0 | 11023 (18787) | 85 |
| | 97.5 | 19006 (29766) | 115 |
| Weibull | 95.0 | 6081 (8219) | 84 |
| | 97.5 | 8193 (10192) | 103 |

**Table S2. Pruning thresholds used for transmission tree sensitivity analyses.** We used a single probability distribution for the distance kernel when assigning progenitors to owned animals and the convolution of the distance kernel for animals for which the owner was unknown and wildlife.

Estimates of introduced cases, with biters directly attributed from contact tracing, are shown in Figure S4A. Between 90 - 400 cases were assigned as introductions (6 - 30 introduced cases per year) over the 14 years depending on the distributions used, with the exception of trees constructed using the Lognormal distribution that linked the majority of cases together under the 97.5% threshold, suggesting that these were too conservative to differentiate introductions from local biters. We generally found good agreement with the contact tracing, assigning around 65% of known biters correctly irrespective of the serial interval and dispersal kernel distributions used. Known biters were rarely mis-assigned as introduced cases, but the probability of misassignment increased at lower pruning cut-offs, with a maximum of ~ 3% of cases mis-assigned using the 95% cutoff with Gamma distributed dispersal (Figure S4B).

We compared the probabilities of biters determined from the consensus tree (Figure S5A), and the resulting distributions of serial intervals and dispersal distances for the probabilistically assigned biter-bitten pairs i.e. excluding traced biter-bitten pairs (Figure S5B). For most cases without a traced transmission link, reasonably plausible biters are identified, and the distributions of serial intervals and dispersal distances align well with the fitted distributions, suggesting that biters are likely detected for a large proportion of cases (see S8). Progenitors were less likely to be assigned correctly within local outbreak clusters, given the plausibility of multiple possible biters (Figure S5A). As expected, using higher pruning cutoffs resulted in longer tailed distributions (more rarer events), and linked cases that were further apart (e.g. beyond 6km). Progenitors that were assigned probabilistically were likely to be further apart than those that were directly traced (Figure S5C), potentially illustrating how contact tracing was easier for transmission events that occurred within a restricted spatial neighbourhood (especially those from the same household).

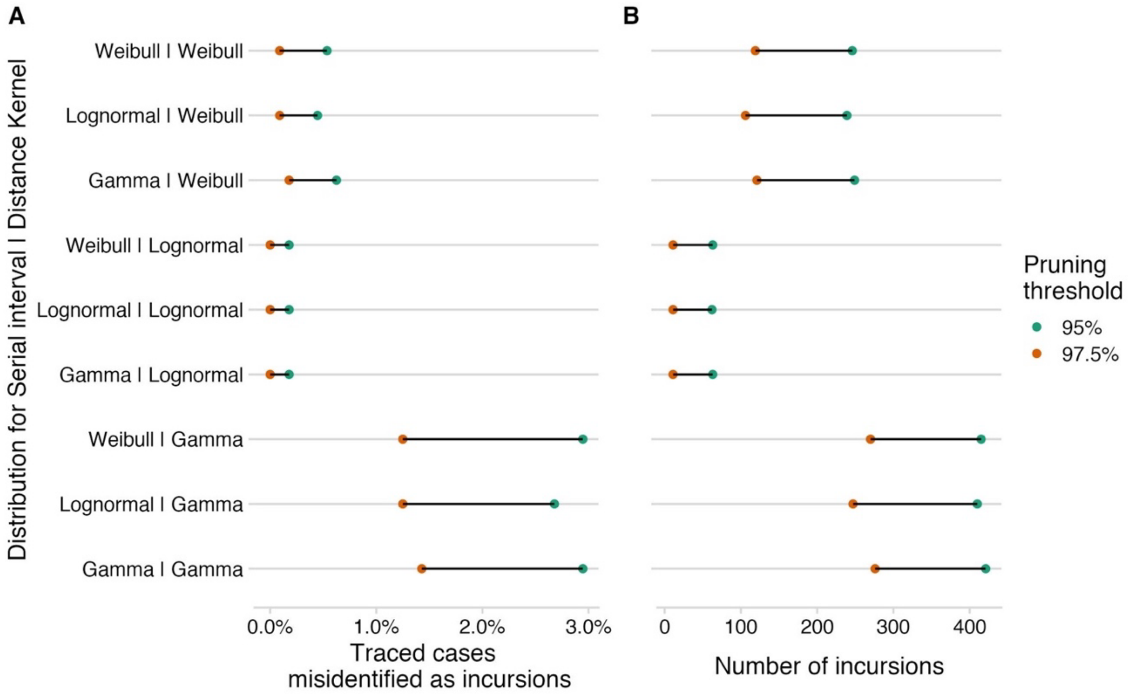


**Fig. S4. Performance of tree-building algorithms for identifying introductions.** (A) The percentage of traced cases misidentified as introductions and (B) the number of introductions generated under the different tree-building algorithms and reference distributions. The y-axis indicates the combination of distributions used for the serial interval and distance kernel, respectively, and the color indicates the pruning threshold.

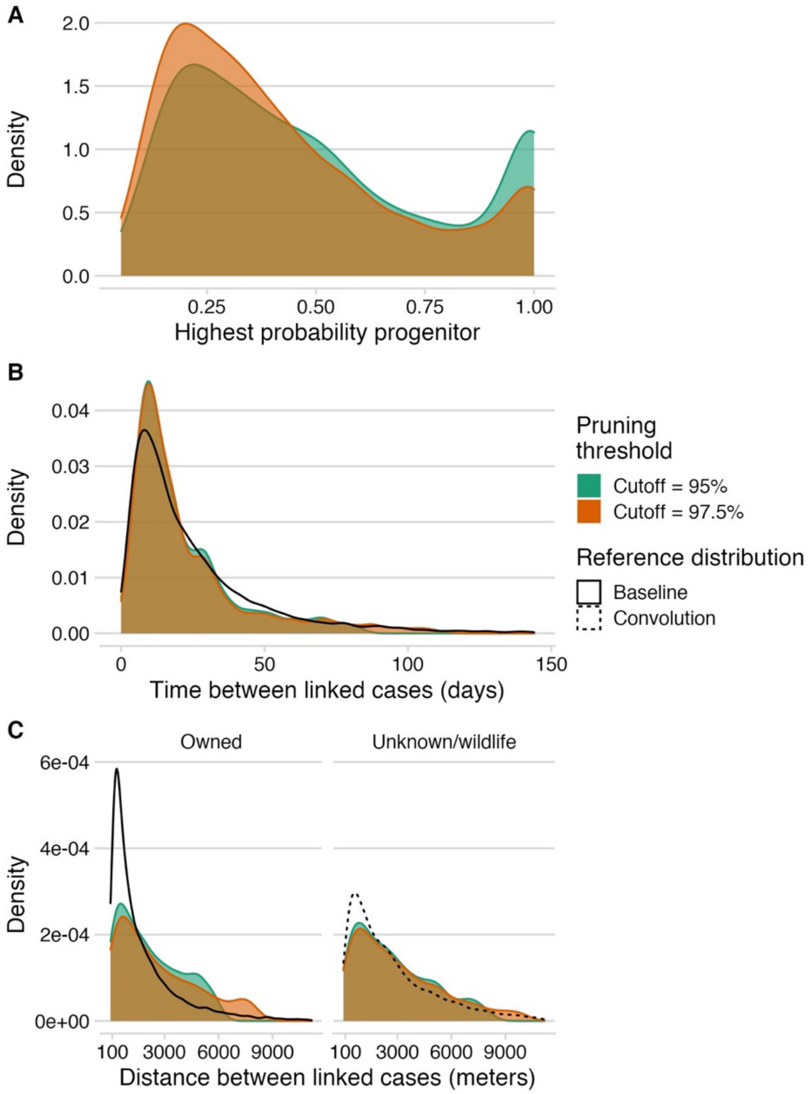


**Fig. S5. Probabilities of inferred progenitors, serial intervals and dispersal distances.** The distributions of (A) the frequencies of the consensus biters (i.e. how often a link was selected from 1000 bootstrapped trees); (B) the serial intervals and (C) distances between the consensus biters and linked cases, excluding cases with a known biter from the tracing data. Left and right panels indicate the pruning cutoff used. The lines show the fitted distributions for the serial interval and dispersal kernel used for the probabilistic transmission tree reconstructions.

From the reconstructed transmission trees we examined clade size (numbers of cases within clades), clade persistence (in weeks), and secondary case distributions (for all cases within the tree). These metrics were largely consistent over the pruning cutoffs, with the lower threshold (95%) breaking apart large clades (shorter tailed distributions compared to the 97.5% cutoff) and generating more singletons (i.e. introductions not linked to any other cases) (Figure S6). To look at uncertainty within the bootstrapped trees, we compared a random sample of 100 trees, to the consensus tree (the set of biters most frequently selected for each case from the 1000 bootstrapped trees), the majority tree (the tree within the bootstrap that had the highest number of consensus biters), and the maximum clade credibility (MCC) tree (the tree within the bootstrap that had the highest overall likelihood, i.e. the highest product of progenitor probabilities). Within tree variation was generally low, but the consensus tree shifted the distribution of distances and times between assigned biter-bitten pairs, because it selects the most likely biter for every case, and therefore tends preferentially to not capture lower probability (rarer) links (Figure S7).

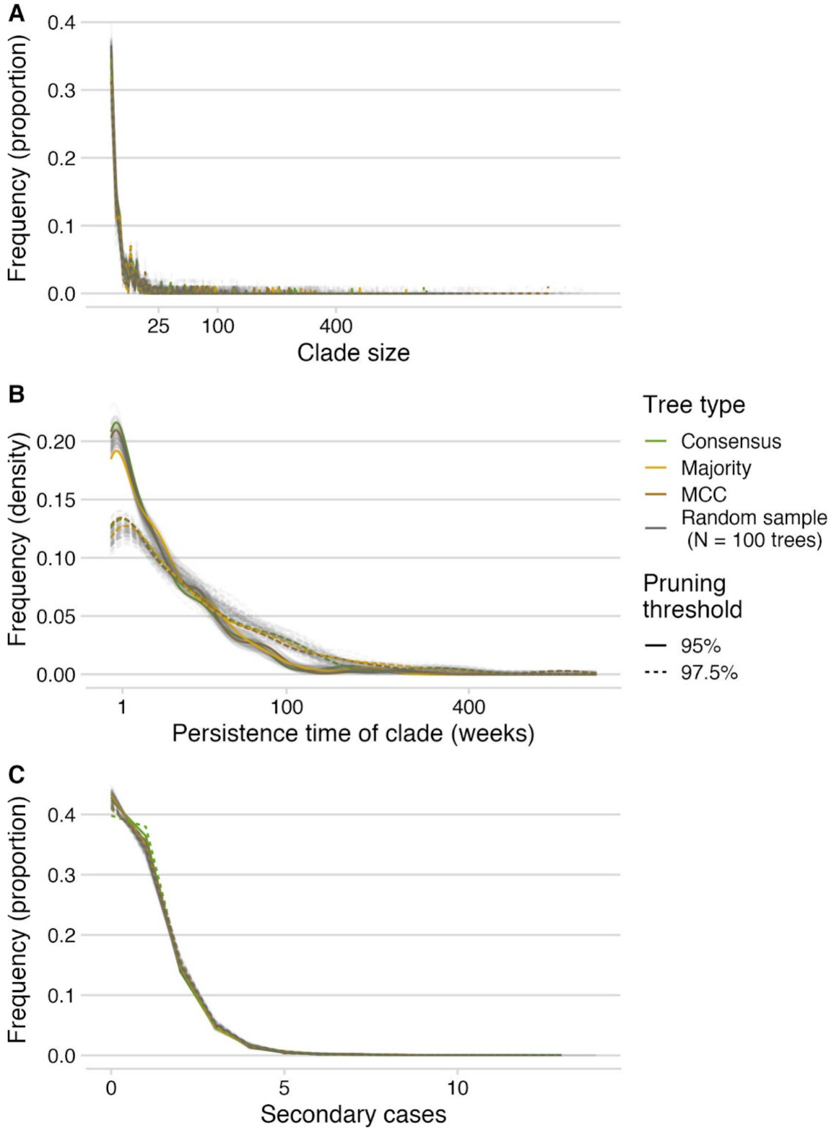


**Fig. S6. Clade metrics under different pruning cutoffs and transmission trees.** Comparing the distribution of (A) clade sizes (x-axis is square-root transformed); (B) clade persistence times (x-axis is square-root transformed), and (C) secondary cases, for a random sample of 100 trees from the bootstrap, the consensus tree, the majority tree, and the maximum clade credibility (MCC) tree.

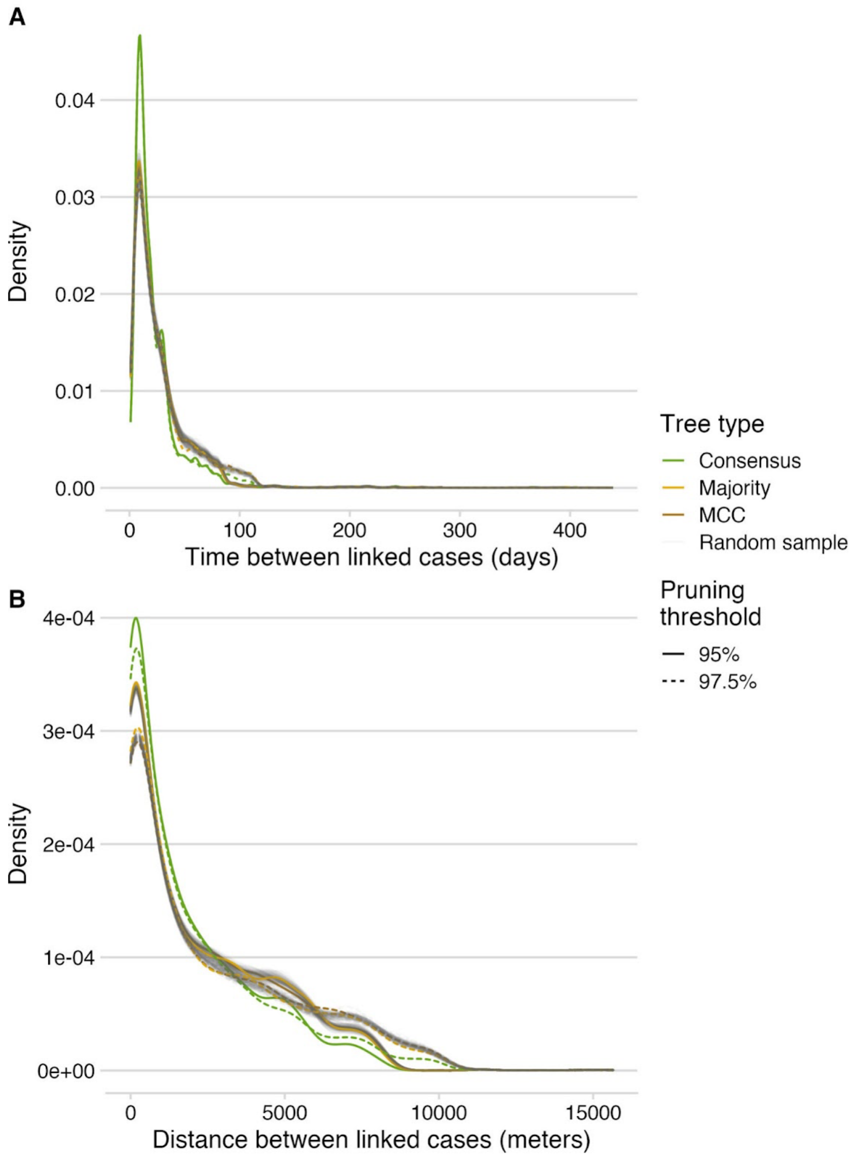


**Fig. S7. Serial intervals and distances between linked cases.** Comparison of the distributions of (A) serial intervals and (B) distances between linked cases for a random sample of 100 trees from the bootstrap, the consensus tree, the majority tree, and the maximum clade credibility (MCC) tree, under the different pruning cutoffs

### The simulation platform

We developed a simulation platform to estimate (using ABC) three contact parameters derived from the Holling functional response, allowing us to determine the functional form of contact in relation to population density: the (inverse of the) search rate, the handling time and the variability in the handling time. We aimed to determine the best spatial scale for modeling rabies transmission by fitting these contact parameters at different spatial scales. Specifically, we assumed that rabid dogs contact other dogs by moving according to the step length distribution fitted from contract tracing (see *Epidemiological parameter estimation* above). At each step, a dog moves within a grid cell or to a new cell, with complete mixing assumed to occur within cells, the size of which is set by the scale of the simulation. Dog population dynamics were modelled to reflect the increasing human population in the district over this time period, incorporating demographic stochasticity and the effect of population turnover on vaccination coverage and susceptible depletion. For modelling dog demography we used parameters akin to an ecological carrying capacity, with increased dog birth and acquisition rates when dog numbers were reduced (through stochasticity or depletion due to rabies) to below carrying capacity.

We programmed the simulation platform in Java. The platform consisted of two layers: a demographic layer modelling dog demographic processes and vaccination campaigns on a regular lattice structure; and an epidemiological layer modelling rabies cases and transmission. The demographic layer tracked counts per grid cell of individual animals by epidemiological status (susceptible, exposed, vaccinated), whereas the epidemiological layer explicitly modelled exposed and infectious individuals in continuous space. All processes were simulated in continuous time. Vaccination campaigns (numbers of dogs per grid cell vaccinated on a pre-specified day), carrying capacity updates and introductions (i.e. cases that originated from outside of the district, either from neighbouring villages or from long-distance human-mediated movement of dogs that were incubating infection) were read in at initialisation. Stochastic demographic events and epidemiological transitions were simulated on-the-fly. This allowed us to combine the simulation of demographic processes using a next-event Gillespie algorithm with exponentially distributed waiting times, alongside pre-computed fixed-time vaccination and introduced cases.

Each simulation run was initialised by reading in: (1) a shapefile of the polygons representing each cell in the grid structure at the given spatial scale; (2) the corresponding carrying capacities per grid cell for each month of the simulation; (3) information on the introduced cases (x-y location, day generated, serial interval, plus auxiliary information) as inferred from transmission trees; (5) the vaccination schedule (grid cell identifier, day on which vaccination took place and number of dogs vaccinated); and (6) static parameters and parameters to be estimated by ABC.

*Demographic and vaccination layer*

We used a grid-based approach, simulating the dog population, N, in each grid cell at the relevant spatial scale, and tracking the number of susceptible, exposed and vaccinated dogs in each cell. Except when rabid (i.e. infectious), dogs were assumed to die at the same per capita rate of 1/25.8 months (6), irrespective of epidemiological status and dog density in the grid cell. Measured and projected dog population sizes were considered to represent carrying capacities K per grid cell (Figure S2). These were pre-computed and updated monthly, with dog demography simulated in the following way. Below the carrying capacity for the grid cell, the rate at which new dogs were acquired or born r was simulated as the maximum of: (a) an acquisition rate of 1/(4.3*30)*K (such that 1/2 of all dogs that died were replaced within 3 months); and (b) the natural grid cell birth rate b*N (where b represents the natural per capita birth rate, set equal to the death rate). This approach was chosen to model a constant acquisition rate by humans per dog below the carrying capacity, until there were sufficient dogs that the local natural birth rate exceeded the acquisition rate. Above the carrying capacity, no additional acquisitions were assumed and births occurred in a density-dependent manner (i.e. r = bN(1 - (N-K)/K)); above 2*K, neither births nor acquisitions were assumed to occur. The number of non-rabid dogs moving between grid cells was assumed to balance out and was not explicitly simulated.

Dog vaccination was implemented according to data on vaccination campaigns (S2 and Figure S3). We used the projected gridded dog population data, together with the data on dog vaccination campaigns by village and the explicit spatial distribution of vaccination status of dogs recorded from the dog population census. We allocated vaccine doses associated with each campaign to cells in the relevant village at the finest ($0.25km^2$) spatial scale, proportionally to the percentage of dogs in that cell recorded as vaccinated. Starting from the finest scale, doses were reallocated to grid cells at larger spatial scales through grid cell aggregation. We assumed vaccine doses delivered at each campaign were first allocated to surviving dogs that had been vaccinated previously, then to new dogs. This conservative approach to vaccination, that assumed preferential repeat vaccination of previously vaccinated dogs, is in line with our experience during vaccination campaigns. Remaining doses were randomly allocated to susceptible dogs across the grid cell. Vaccine protective efficacy was assumed to be 100%.

*Epidemiological layer*
The epidemiological layer was individual-based, with basic accountancy applied to the demographic layer at each epidemiological transition to ensure the consistency of the number of susceptible and exposed individuals in the relevant grid cell. When transmission occurred, a new case record was created for the exposed individual and an infectious period and serial interval (combining incubation and infectious periods) sampled jointly from the data. On reaching the date of onset of clinical signs, movement and biting were simulated.

Movement and biting for each focal individual were simulated for the duration of its infectious period in the form of a sequence of steps, a process which continued until the end of the infectious period when the dog was assumed to die. These steps constituted encounters that could potentially lead to transmission at a particular location in space, and were drawn from the step length distribution fitted from the contact tracing data (see *Epidemiological Parameter Estimation* above). These steps were modelled following a random walk in continuous space, with the contact rate within the grid cell for each step modelled according to a Holling Type II functional response as explained below.

To simulate contact, a time duration was drawn for each step from an exponential distribution with rate parameter rc,d (see below for details). At the end of each step (time period), assuming that the grid cell was not empty and that the infectious period had not ended, a bite was simulated. For each bite, a dog was selected with equal probability among those in the grid cell (equivalent to complete mixing within the gridcell), and a transmission event occurred with probability 0.49 (1) if the bitten dog was susceptible (i.e. neither vaccinated nor already incubating infection). We kept track of dogs that had been exposed previously, allowing us to ensure that each dog was only infected once. Demographic processes were temporarily interrupted while a case was infectious, implying that in relation to vaccination and demographic processes, all secondary infections occurred simultaneously. This simplification, justified by the short infectious periods of rabid dogs, was made in the interests of computational efficiency and algorithmic simplicity. If a step moved a dog to outside the modelled area (occupied cells and a

1km buffer beyond these), further steps were simulated until obtaining one within the modelled area, effectively allowing rabid dogs to rebound into populated areas and/or the surrounding buffer. These may have acted like introductions into the district, slightly affecting some of our estimates, but because of the high probability of rebounding into the buffer area where onward transmission was not possible, we do not expect these to have had a major influence on our results.

We did not find a significant relationship between step lengths and population density from the contact tracing data, so we assumed in the simulation that all steps were independent and identically distributed. We therefore applied uniform random bearings and step lengths drawn from the fitted Weibull distribution (Figure S1C) corresponding to the steps taken during an animal's infectious period. To capture human-mediated movement, we implemented the first step per dog as long-distance transport with probability 0.02, assuming low frequency human-mediated transport (7), to a randomly selected occupied cell chosen with probability proportional to K. We applied this probability only to the dog's first step assuming that most long distance transport occurs only once, typically during the dog's incubation period.

The expected contact rate $r_{c,d}$ in a grid cell was modelled as a Holling Type II functional response, formulated as a function of dog density in the cell and parameterised by dog-specific handling and search rate parameters. Specifically, in a given grid cell, the expected daily contact rate was computed as:

$$r_{c,d} = ([\delta+1]\, r_{s,d}) / (1 + (r_{s,d}\, T_{h,d} * [\delta+1]))$$

where dog density δ is in units of dogs/km$^2$ (computed at a given spatial scale), the dog-specific handling time $T_{h,d}$ captures the mean time spent with each dog encountered, and the search rate $r_{s,d}$ the rate at which the focal dog made contact per unit dog density, capturing search efficiency. Note that in a 1km$^2$ grid cell, density is equal to the number of dogs per cell, and the inverse of the search rate $1/r_{s,d}$ can be interpreted as the expected time until an encounter with a specific dog within the cell. We chose to add one to the density to avoid generating an encounter rate of zero in empty cells; this allowed infectious dogs to move on from empty cells after a finite time.

To model individual variability, the functional response was allowed to vary between dogs. That is, for each rabid animal, an expected handling time $T_{h,d}$ was drawn from a Gamma distribution, parameterised by mean $T_{h,mean}$ and scale parameter $T_{h,scale}$ estimated via ABC. $T_{h,d}$ was used to compute the search rate ($r_{s,d}$), noting that the ratio of the inverse of the search rate ($1/r_{s,d}$, which can be interpreted as the expected search time) to handling time was held constant for all rabid animals in the simulation. We used this formulation to keep the level of density dependence (measured as the ratio of search time to handling time) constant between dogs while introducing individual variation through the handling times. The dog-specific values $T_{h,d}$ and $r_{s,d}$ were then used to assign the functional response for each dog.

Model Fitting using Approximate Bayesian Computation

Using ABC we estimated three parameters that relate to the contact function: the mean and shape of the handling time ($T_{h,mean}$ and $T_{h,shape}$), and the inverse of the search rate, $1/r_{s,d}$. We simulated 50,000 parameter sets from across the pre-selected parameter range and jointly estimated these three parameters. Parameter sets were drawn uniform randomly from the following ranges in units of days: 0.5-10 for $T_{h,\ shape}$, 0.001-3 for $T_{h,\ mean}$ (the scale parameter $T_{scale}$ for the handling time was then computed as the mean/shape) and 0-25 for $1/r_{s,d}$ (Figure S9). These same parameter sets were used to simulate rabies dynamics at each spatial scale.

For the ABC, we employed three acceptance criteria designed to capture the spatiotemporal characteristics of rabies dynamics based on: 1) the variability in monthly counts of rabies cases; 2) the dog density at which cases occurred (and therefore case clustering by density); as well as 3) biologically plausible limits to rabid dog biting behaviour (numbers of bites per rabid dog). Following a first round of ABC, we simulated from the accepted (first round posterior) parameter sets to determine how reliably these reproduced outcomes that were consistent with the ABC criteria, and used reliability to define a final posterior distribution and establish the best spatial scale for modeling rabies transmission. We developed the ABC criteria from the full dataset of carnivore cases (n=3,281), i.e. domestic dogs, cats, and wildlife, assuming that livestock do not contribute to onward transmission.

Criterion 1 was designed to capture key characteristics of the observed time series, and was based on an ABC measure of distance that included outbreak sizes but not necessarily timing. Specifically, we compared the distribution of the number of cases per month between the data and simulation runs. To generate a distance metric, we aggregated cases per calendar month and used the fitdistrplus package in R to fit a Negative Binomial distribution to these data by maximum likelihood. We simulated 100,000 draws from this fitted distribution to generate a discrete distribution with breaks every 5 cases (Figure S8A, grey shaded area) and computed pointwise 99.9 percentile intervals. Parameter sets were accepted if no more than two points from the simulation run fell outside these computed bounds.

Criterion 2 was designed to capture the observation that cases tended to occur in areas of higher than average dog density (Figure 1D). We therefore used a measure of the ABC distance between the data and simulation outputs in the distribution of dog densities at which cases occurred. By design, the simulator was not able to generate cases in areas of zero dog density, so we first moved cases from the data occurring at such locations (corresponding to locations of unknown dogs found dead, n = 93) to the closest occupied cell. Because the distribution of case location densities was highly skewed, we log-transformed the data using the transform log(density + 0.1), to generate a more symmetric distribution, relative to which deviations between the data and simulation runs would be more evident. Dog populations increased considerably over the period, therefore it was important to standardise dog population densities at which cases occurred relative to the contemporaneous distribution of dog densities. To achieve this, a z-score was computed on the logged population densities. For each case, we computed a time-consistent standardised log density (sld) at the time when the case occurred as:

$$\mathrm{sld}_c = [d_c - \mathrm{mean}(d_{t,g})] / \mathrm{sd}(d_{t,g})$$

where $d_c$ denotes the measured log-transformed density at the location of the case, mean($d_{t,g}$) is the mean of the log-transformed densities of all grid cells at the time $t$ of the case occurrence, and sd($d_{t,g}$) the standard deviation. Aggregating sld$_c$ values at intervals of 0.25 produced a discrete distribution (Figure S8B, red dots). We generated intervals using a bootstrapping procedure according to which the observed number of rabid carnivore cases (3,281) were sampled with replacement from the data 100,000 times, their sld values aggregated, and a pointwise range computed (Figure S8B, grey shading). Parameter sets were accepted if no more than four points from the simulation run fell outside these computed bounds.

Criterion 3 was a cap on rabid dog biting behaviour, assuming that rabid dogs never bite more than 100 other dogs during their infectious period i.e. runs that generated dogs that bit >100 other dogs were rejected under the ABC. The most prolific biter in the contact tracing data was observed to bite 72 other dogs, therefore the cap of 100 allows for some under-detection.

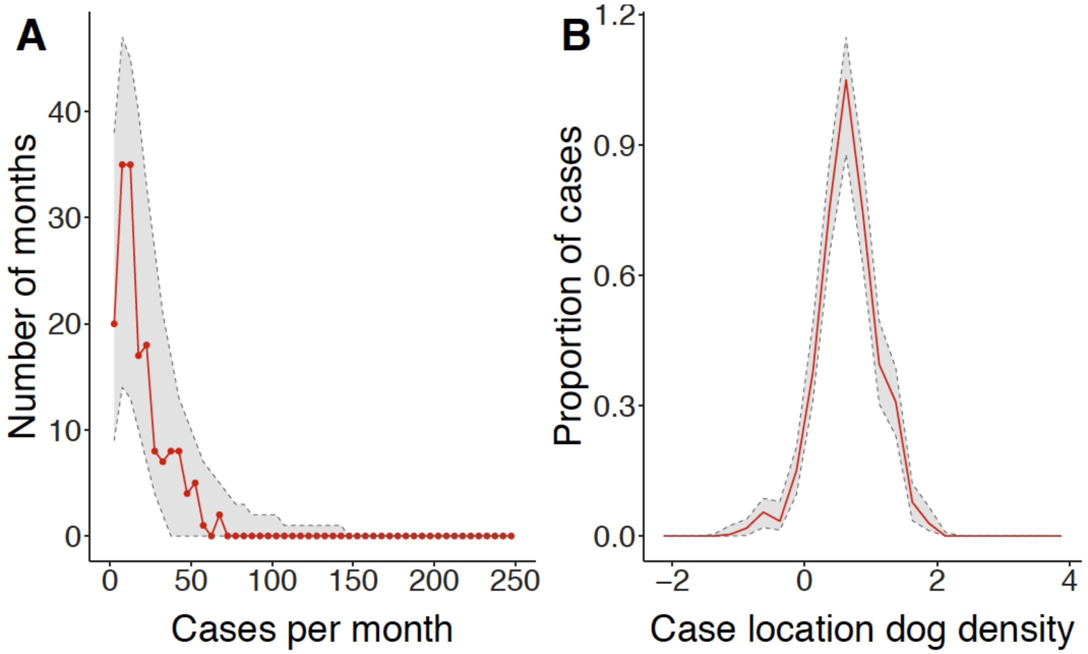


**Fig. S8. ABC criteria for model fitting.** (A) Cases per month and (B) Normalised dog densities at which cases occurred. The data are shown in red and bootstrapping was used to compute the bounds of these distributions (grey shading), as described in S6.

The ABC criteria excluded the vast majority of parameter sets across all spatial scales (Figure S9), with the highest number accepted at the 1x1km scale (N=168). Accepted parameter sets were generally those associated with low levels of density dependence. At larger spatial scales, parameter estimates were subject to greater uncertainty, especially in the search rate. Although the values for the shape parameter of the handling time spanned the range 2.784-9.927, the bottom of the range that makes the largest qualitative difference to the shape of the gamma distribution was well-identified, and given with the mean of the handling time, all of the parameter sets in the posterior generated gamma distributions with a qualitatively similar shape with minimal skew and similar variance.

We re-simulated rabies dynamics from the posterior distribution of accepted parameter sets following the first round of ABC at each spatial scale, simulating 100 runs per parameter set, to evaluate how reliably they reproduced the data. On re-simulating from initially accepted parameter sets we found that the most reliable parameter sets were at the 1km2 scale (Figure S10), where maximum reliability exceeded 60%. Parameter sets with at least 50% of runs meeting the ABC criteria were considered sufficiently reliable and retained within the final posterior distribution (ranges for: the mean of the handling time Th,mean 0.572-0.781, the shape parameter of the handling time Th,shape 2.784-9.927, and the mean search time 1/rs,d 0.056-1.541). Overall, our findings are consistent with the suggestion that the apparent relationship between density and transmission depends on the spatial scale of measurement (8).

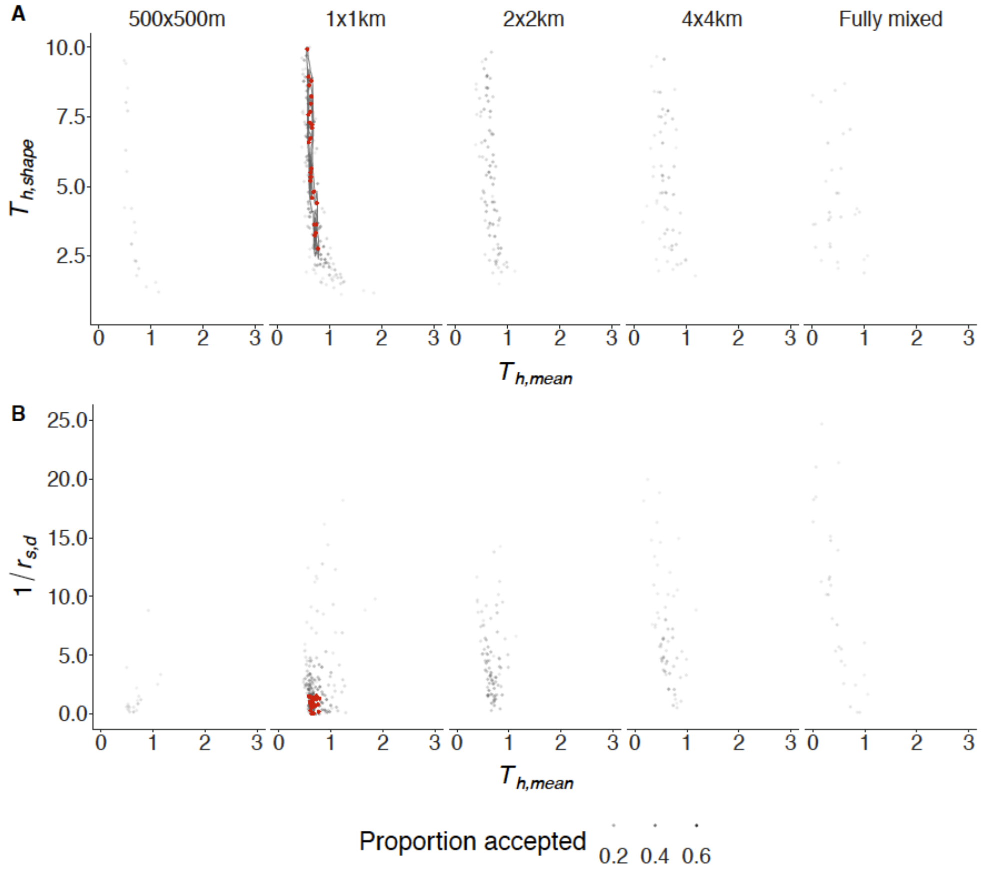


**Fig. S9. Posterior distributions for parameter combinations at each spatial scale.** All parameter sets accepted in the first round of ABC are shown, with shading denoting the proportion accepted in the second round of ABC, and final posterior parameter sets (proportion of the 100 runs accepted in the second round of ABC exceeding 50%) shown in red, with contours highlighting density in the final posterior.

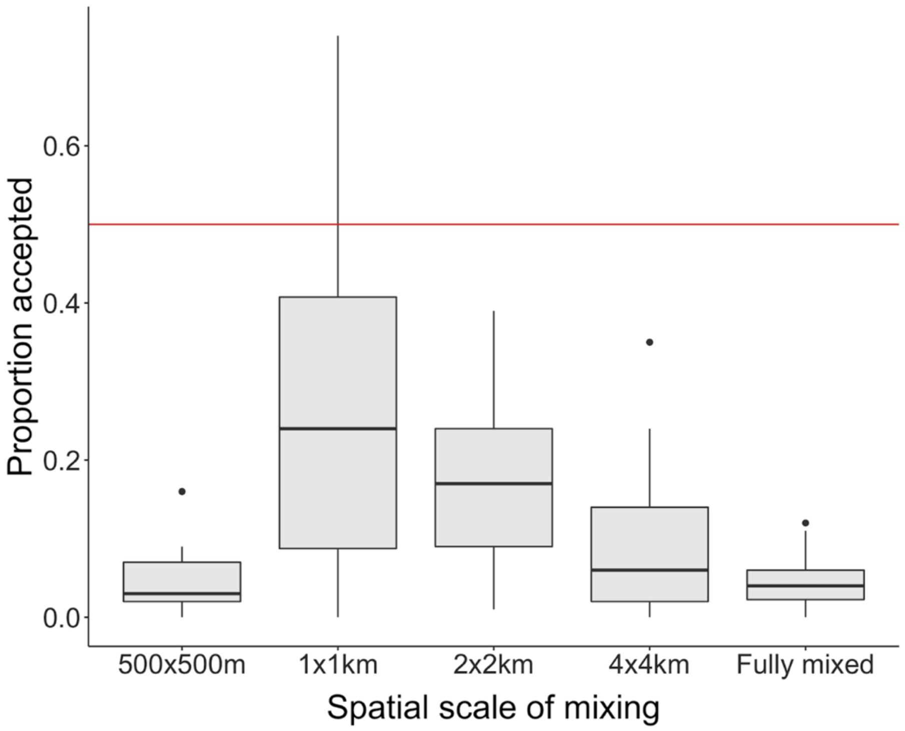


**Fig. S10. Reliability of parameters sets at each spatial scale.** Boxplots show the interquartile range, 95 percentiles and outliers from 100 simulations of each initially accepted set of parameters at each spatial scale. The red line indicates the reliability threshold above which we selected the final posterior distribution from the 1x1km scale. At this scale the ranges for the mean of the handling time $T_{h,mean}$ was 0.572-0.781, the shape parameter of the handling time $T_{h,shape}$ was 2.784-9.927, and the mean search time $1/r_{s,d}$ was 0.056-1.541.

Simulations from the posterior distribution and scenario exploration

We simulated from the final posterior distribution to explore counterfactuals and understand how different components of transmission contribute to epidemiological dynamics. We examined vaccination impacts by simulating with vaccination campaigns as reported in the data, and with reduced levels of vaccination consistent with those in nearby districts. We examined the dynamic consequences of individual-level heterogeneity by comparing our simulations to those where we constrained individual variation in contact (i.e. setting the handling time for all dogs as equal to the mean so that all dogs had the same parameter values). We investigated how clustering influenced the basic and effective reproductive numbers ($R_0$ and R) by treating cases generated from the posterior simulations as index cases and comparing the resulting secondary case distributions (i.e. with and without constraints due to clustering and susceptible depletion), examining $R_0$ and R over time and across space (i.e. by 1km$^2$ grid cell). We also investigated the contribution of introductions to persistence by simulating epidemics with introductions occurring throughout (and vaccination campaigns as per the data) and with introductions occurring only during the first 12 months of the simulations, and comparing these with the transmission trees inferred from the contact tracing data.

In general, simulated levels of rabies incidence were skewed, with a lower median than mean incidence over the 14 years, and a long tail of high incidence realizations. When vaccination was omitted entirely, some extreme outbreaks resulted that we considered to be unrealistic but under the low vaccination scenario just 1% of runs exceeded 50,000 cases which was the criteria we used to end unrealistic simulations. We suspect that in practice these extremes do not occur because they are mitigated by localized behavioural changes like reactive vaccination, culling and restrictions to dog movement that were beyond the scope of our model (further analyses of the contact tracing data show that previous rabies cases and human rabies exposures in the previous month increase the likelihood that a rabid dog will be killed or tied before biting). Simulations with very low levels of vaccination (~7%, equivalent to levels in neighbouring districts) led to increased numbers of cases (Figure 3B versus A) and a stronger shift in case occurrence towards higher density areas (Figure S12). The median case counts under observed coverage were over three times lower than simulated case counts under low coverage. Using the ratio of exposures and deaths per case from the data (0.45 exposures per case and 0.013 death per case), we also estimated human rabies exposures and deaths prevented through mass dog vaccination.

In simulations without individual variation in dog contact parameters, rabies incidence was reduced by almost 50% and large outbreaks did not occur (Figure 3C versus A). With introductions inferred from the transmission trees used to seed outbreaks over the full 14 years, 100% of simulations lasted the entire duration (i.e. from 2002-2015). However, if introductions were restricted to just the first 1 year of the simulations, the median duration of outbreaks was just over two years (27 months), and rabies persisted for less than four years in over 90% of simulations, with the longest simulation lasting 8 years (Figure 3A versus D).

We found that frequency distributions of clade sizes, persistence times and secondary cases were superficially similar for the simulations (under observed vaccination campaigns and for scenarios with minimal vaccination, N = 100 subsampled simulations from the posterior for each scenario) and the reconstructed transmission trees (Figure S11A-C, N = 100 random subsampled trees from the bootstrap, in addition to the majority, MCC, and the consensus trees). But there were evident differences when looking at specific characteristics of these distributions (mean, median, and maximum, Figure S11D-F). Simulations with vaccination as per the data tended to produce smaller clades on average than reconstructed transmission clades that persisted for shorter lengths of time. This is consistent with our estimates of underdetection, suggesting that contact tracing is more likely to miss introductions that result in deadends than cases that cause secondary transmission. Likewise the algorithm with pruning may underestimate the number of clades and overestimate their persistence by linking unrelated co-circulating clades. Without vaccination, the maximum of all the metrics shifted upwards, and some clades persisted for the duration (14 years). However, mean simulated clade duration was between 1-2 years in the absence of vaccination.

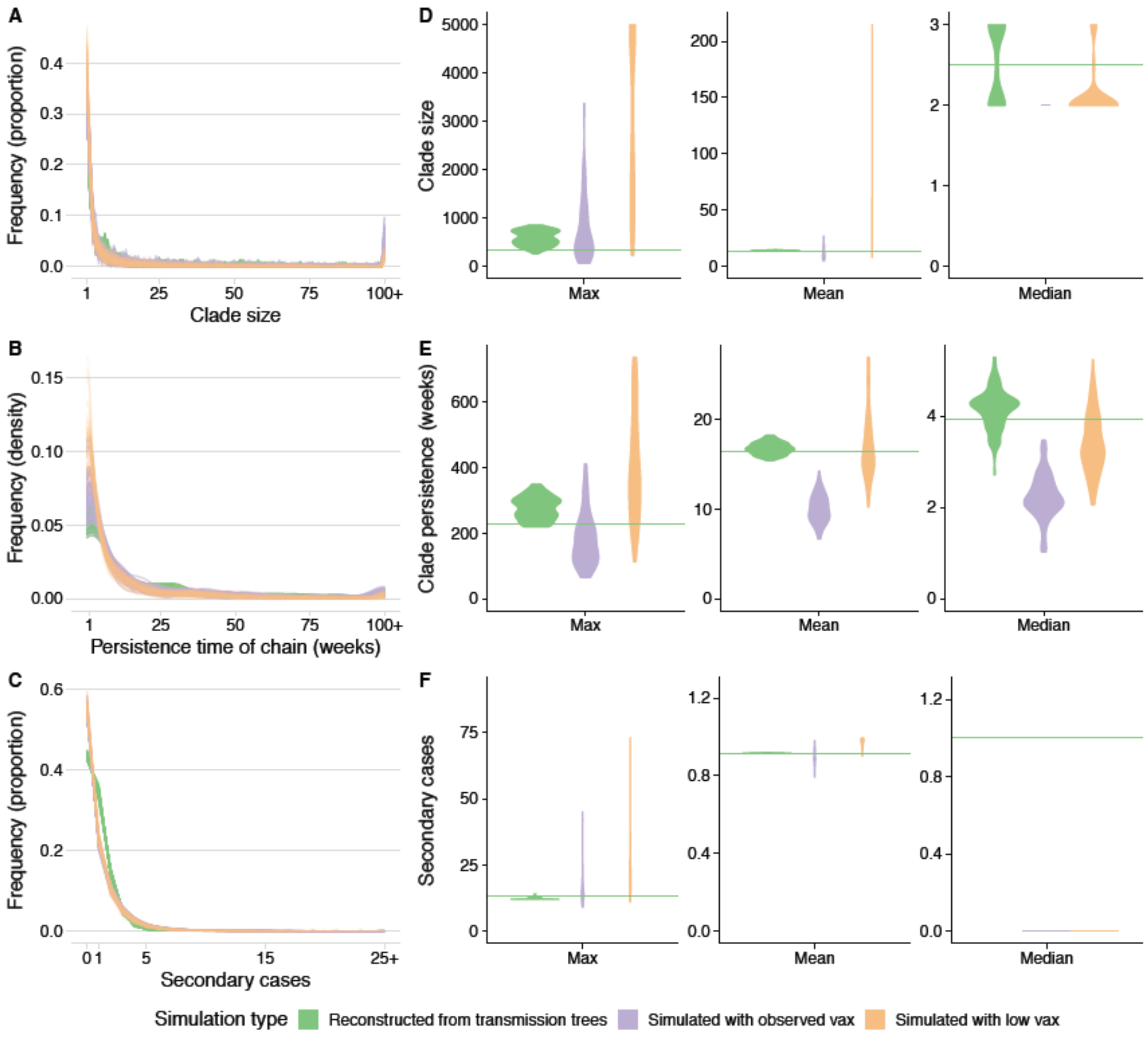


**Fig. S11. Comparison of metrics from transmission trees and simulations.** Distribution of (A) clade sizes (B) clade persistence times and (C) secondary cases for a subsample of transmission trees and for simulations from the posterior distribution. For the transmission trees we used the 95% pruning cutoff (N = 103, including a random subsample of 100 trees, as well as the consensus tree, the MCC tree, and the majority tree) and for the simulations from the final posterior distribution we modelled vaccination as observed (purple) and with minimal vaccination (orange), and plotted the distributions from 100 runs of each scenario. Values greater than 100 for (A) and (B) and 25 for (C) are grouped together. The corresponding max, mean, and median of (D) clade sizes, (E) clade persistence times and (F) secondary cases for these trees are also shown. In (D) to (F) the green line shows the corresponding value for the consensus tree. The maximum clade size under low vaccination coverage exceeded 15,000 cases so we censored the figure at 5,000.

We calculated $R_0$ by randomly selecting dogs from across the landscape in proportion to dog density, and using these as index cases from which to simulate secondary cases (Figure S12). When selecting dogs at random (and therefore in proportion to dog density) this approach led to an $R_0$ of 1.47 (95% Percentile Intervals (PI) 1.39-1.56) consistent with previous estimates from this population using different analytical approaches (1). Selecting dogs that were infected with rabies in our endemic simulations as index cases resulted in a higher $R_0$ estimate of 1.48 (95% PI 1.38-1.58), whereas randomly selecting index cases by location (grid cell) led to a slightly lower estimate (1.35, 95% PI 1.27-1.43). These simulations illustrate how the spatial location of cases and local dog density alters transmission potential and therefore how $R_0$ is an emergent property of localized patterns of circulation within heterogeneously distributed populations. Even without strong density-dependent effects on contact rates, the tendency for infection to circulate longer and infect more individuals in higher density areas increases $R_0$.

We investigated how spatial clustering of rabies cases and of heterogeneity in dog density influenced transmission by treating simulated cases under the endemic scenario (with low vaccination coverage) as index cases, ie. without any effects of susceptible depletion, and comparing dogs bitten and secondary cases per case, across 1000 simulations. Under the minimal vaccination coverage scenario, rabid dogs bit on average 2.23 other dogs (95 percentile intervals (95%PI): 2.08-2.32), comprising 0.164 vaccinated dogs (95% PI: 0.131-0.202), 0.152 dogs incubating infection (95% PI: 0.113-0.193) and 1.912 susceptible dogs (95% PI: 1.81-1.96) leading to an R of 0.983 (95% PI: 0.919-1.009). In contrast, on removing the effects of local depletion, the corresponding index cases bit on average 2.92 susceptible dogs (95% PI: 2.74-3.07) leading to R of 1.48 (95% PI: 1.38-1.58). The difference in R between these scenarios was due to vaccination (7.4% of bitten dogs were vaccinated in the low vaccination endemic circulation scenario), prior exposure (6.8% of bitten dogs were already incubating infection) and reduced contact opportunities due to prior rabies deaths (23.3% fewer contacts per index case), which together resulted in about one less effective (susceptible) contact per rabid dog. Discounting the effects of vaccination, R was reduced by 30%, of which 78% was from reduced numbers of susceptible dogs, and 22% due to re-exposure of dogs already incubating infection.

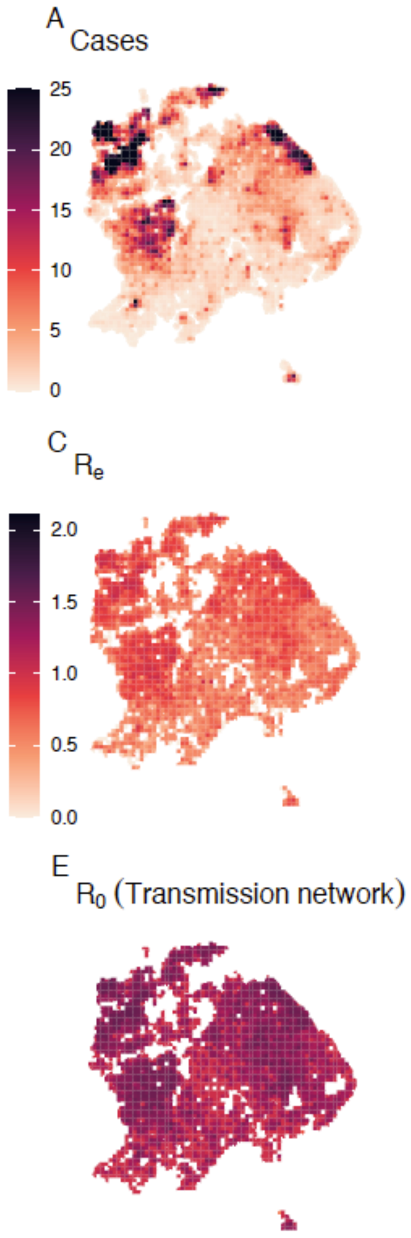


**Fig. S12. The spatial distribution and variability in rabies transmission.** Maps illustrate (A) simulated spatial variation in rabies cases under a scenario of low vaccination coverage; (B) dog density at the midpoint of the timeseries; (C) the simulated effective reproductive number R, under the scenario of low vaccination coverage (i.e. based on realized cases from the simulation shown in A); (D) $R_0$ from simulated index dogs selected in proportion to dog density, i.e., as per B (with no susceptible depletion effects); (E) $R_0$ from simulated index dogs selected from cases from the endemic transmission network under the scenario of low vaccination coverage (scenario A, with susceptible depletion effects removed) and (E) $R_0$ from simulated index dogs selected by grid cell.

Estimating case detection
Contact tracing undoubtedly results in underdetection of some rabies cases, but methods to estimate detection probability are limited. Since the vast majority of observed cases could be linked to plausible progenitors, we believe that our contact tracing detected a high proportion of cases. This was backed up through exploration of the tree-building algorithm's performance when using a convolution of both the serial interval and dispersal kernel to assign progenitor, which may be better suited to linking cases with unobserved intermediates (results not shown). Working from recently developed analytical methods (5), we estimated the probability of case detection from the distribution of the number of unobserved cases, κ, between observed linked cases inferred from contact tracing data via the transmission tree reconstructions.

We focus on time intervals between cases because this is an additive measure, unlike distance between cases, where linked cases can be separated by many unobserved intermediates but still occur close in space due to sequential host movements. Assuming that all cases have the same probability of being detected $\pi$, the probability of not observing $k$ intermediates between linked cases is given by the geometric distribution:

$$f(\kappa = k) = \pi(1-\pi)^{k-1} \qquad \text{Eq 2}$$

Figure S13A shows this expected distribution across a range of detection probabilities, $\pi$. We simulate $\kappa$ (the number of unobserved intermediates between two observed linked cases), by taking the sum of different numbers of draws ($\sum_{i=1}^{k} x_i\, , k \;=\; 1\,..\,k_{max}$) from the serial interval distribution (Lognormal, Table S1) and picking the value of $k$ that generates the closest fit to the observed time difference between the two observed cases. We then minimize the sum of squared differences (SS in Eq 3) between the simulated normalized distribution of $k$(denoted $p_k$) generated across all observed linked pairs, and those generated by Eq.2 for detection probabilities, $\pi$, ranging between 0.01 and 0.99.

$$SS = \sum_{k=1}^{k_{max}} (p_k \;-\; f(\kappa = k \mid \pi))^2 \qquad \text{Eq 3}$$

To account for the long-tailed serial interval distribution, we adapted this approach, by ranking and matching the first set of simulated intervals from the $k$ draws against the observed intervals. This was to better account for long incubators (that are to be expected in proportion to the number of observed cases), which are, with high probability, assigned to result from unobserved intermediates rather than being directly linked to their biters.

To test the performance of these two methods, we simulated intervals between linked cases given a known detection probability $\pi$, drawing values of $\kappa$ from the distribution defined by Eq 2 and summing the equivalent number of draws from the serial interval distribution for 3000 cases. We then applied the fitting approach described above, generating 10 estimates per simulation for 1000 simulations from Eq 2 across a range of detection probabilities (Figure S13B). We also examine the accuracy of this case detection estimation approach applied to simulations from the final posterior distribution of our individual-based model at the 1x1 km scale. Specifically, we applied a detection probability to ‘observe’ cases from 100 simulated runs with vaccination as observed. We then estimate case detection from the resulting intervals between linked case pairs in this observed set, generating 10 case detection estimates per simulation (Figure S13B).

We find that in general, estimates of case detection are recoverable. The case detection estimation approach using ranked matching performs better than the naive approach, at detection probabilities exceeding 0.75 (Figure S13B), but tends to underestimate true detection probabilities by about 10% at levels of case detection between 0.3 - 0.75.

We apply the ranked matching approach to transmission trees reconstructed with the best fitting parameter distributions (Lognormal for $S$ and Weibull for $K$). As pruning removes links between cases that exceed a probability threshold, detection estimates will likely be sensitive to what we set this threshold to be. We compared three pruning thresholds: 95%, 97.5% and unpruned (i.e. where all observed cases are assigned a progenitor in the dataset) to generate a range of detection estimates. We subsampled 100 bootstrapped trees and also included the majority tree and the maximum clade credibility tree (*Transmission tree reconstructions*). For each tree, we took the mean of 10 estimates as the detection probability.

Overall, we estimate case detection probabilities between 0.83 - 0.95 depending on the pruning threshold (Figure S13C). We also estimate the probability of detecting at least one case in a clade given a clade size: under the assumption that all cases in a clade have the same detection probability ($\pi$, estimated in Figure S14C), the probability of detecting at least one case in a clade of size $N$ is $1 - (1 - \pi)^N$ (or 1 - the probability of not detecting any of the N cases in the clade). Based on our estimates of case detection probabilities, we likely detect almost all clades that have more than four cases (probability > 0.99).

There are many limitations to this approach. These estimates are sensitive to the underlying serial interval and distance kernel distributions used to reconstruct transmission trees, and the transmission tree reconstruction is unlikely to perform well with significant underdetection of cases. In these situations, integrating additional data streams, such as phylogenetic data, may improve estimation and the accuracy of transmission tree reconstructions generally. In addition, the main assumption of these methods is that all cases have the same detection probability, which is unlikely to be realistic across space and over time. Detection probability is likely to correlate with clade size such that small clusters (or single cases) go undetected. While there are methods for estimating the number of undetected importations from clade size data, they rely on strong assumptions about the secondary case distribution and its relationship to clade sizes. Our modelling shows that naive expectations of clade sizes do not capture the impact of local susceptible depletion or transmission heterogeneity. Further work is needed to validate and improve these methods and tackle outstanding gaps in methodology. Nonetheless, these case detection estimates provide some confidence that overall our contact tracing detected the majority of rabies cases.

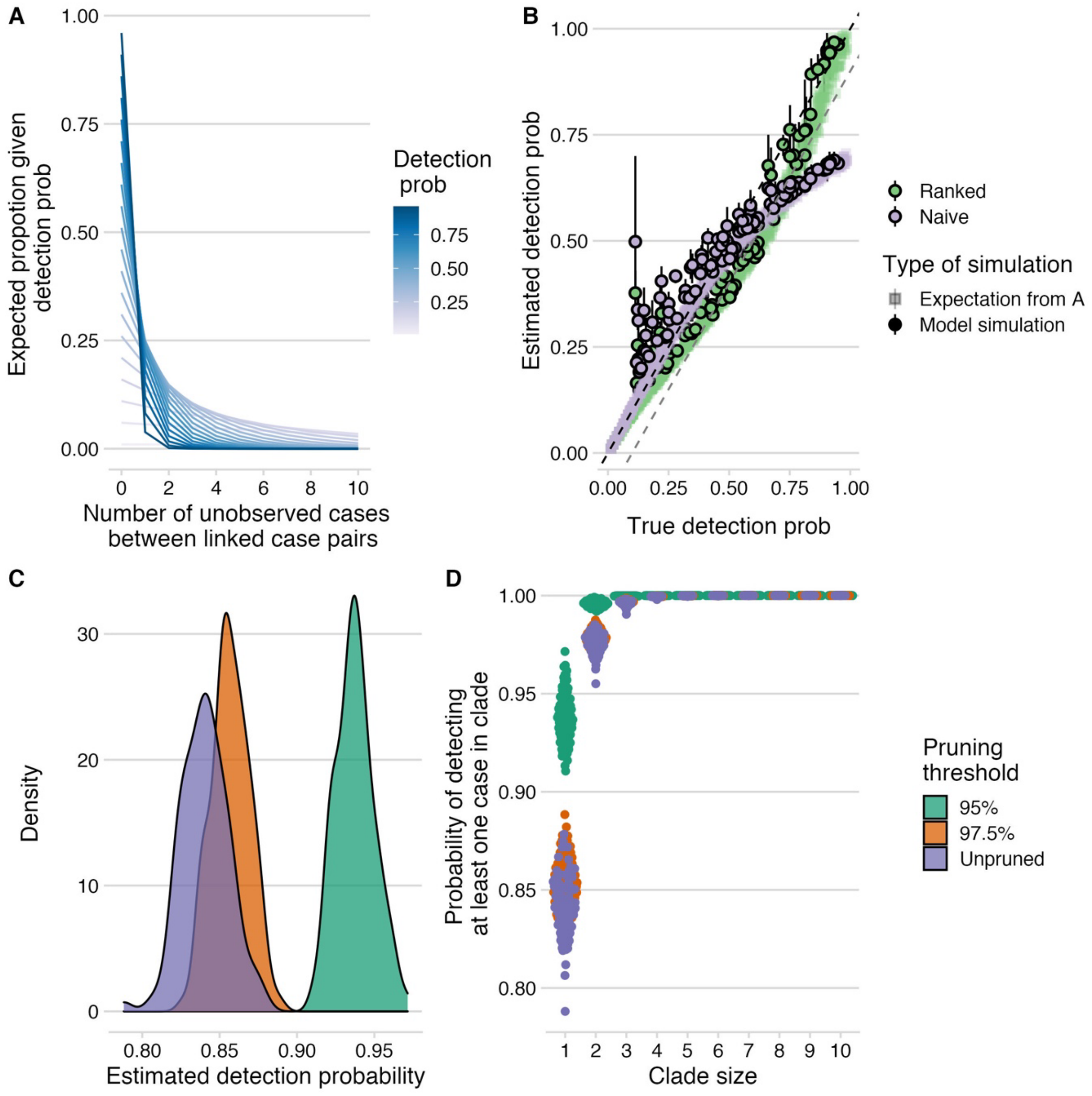


**Fig. S13. Estimation of case detection probabilities.** (A) Expected proportion of linked cases with $\boldsymbol{\kappa}$ unobserved intermediates (x-axis) for a range of case detection probabilities, assuming that detection probability is the same for all cases. (B) Estimated detection probabilities (mean and range) from simulated time intervals between linked cases given a known detection probability (x-axis). Shapes indicate the type of simulation, with squares indicating simulations generated assuming the geometric distribution in A, and circles indicating simulations from the final posterior distribution of our model (N = 100, see S6). Colors indicate whether at the first time step, ranked matching between simulated and observed times was used to capture the long-tail of the serial interval distribution. The black dashed line shows the 1:1 line and the grey dashed line the 1.1:1 line. (C) Detection estimated from times between linked cases from transmission trees (with pruning cutoffs of 95%, 97.5%, and unpruned, i.e. where all cases are assigned a progenitor). (D) Probability of detecting at least one case given estimated detection probabilities and clade sizes (x-axis).
